# Abo1 ATPase facilitates the dissociation of FACT from chromatin

**DOI:** 10.1101/2024.06.17.599424

**Authors:** Juwon Jang, Yujin Kang, Martin Zofall, Carol Cho, Shiv Grewal, Ja Yil Lee, Ji-Joon Song

## Abstract

The histone chaperone FACT is a heterodimeric complex consisting of Spt16 and Pob3, crucial for preserving nucleosome integrity during transcription and DNA replication. Loss of FACT leads to cryptic transcription and heterochromatin defects. FACT was shown to interact with Abo1, an AAA+ family histone chaperone involved in nucleosome dynamics. Depletion of Abo1 causes FACT to stall at transcription start sites (TSS) and mimics FACT mutants, indicating a functional association between Abo1 and FACT. However, the precise role of Abo1 in FACT function remains poorly understood. Here, we reveal that Abo1 directly interacts with FACT and facilitates the dissociation of FACT from chromatin. Specifically, the N-terminal region of Abo1 utilizes its FACT interacting (FIN) helix to bind to the N-terminal domain (NTD) of Spt16. In addition, using single-molecule fluorescence imaging, we discovered that Abo1 facilitates the ATP-dependent dissociation of FACT from nucleosomes. Furthermore, we demonstrate that the interaction between Abo1 and FACT is essential for maintaining heterochromatin in fission yeast. In summary, our findings suggest that Abo1 regulates FACT turnover in an ATP-dependent manner, proposing a model of histone chaperone recycling driven by inter-chaperone interactions.

## Introduction

The genomic DNA of eukaryotic cells is packed in the nucleus in the form of chromatin along with histones. The organization of chromatin is highly regulated by various histone chaperones which facilitate chromatin dynamics and gene expression (1). Chromatin is assembled stepwise by histone chaperones (2). Two copies of H3-H4 come first on the DNA creating a tetrasome, and two copies of H2A-H2B are sequentially assembled into the tetrasome, creating a hexasome and a nucleosome (3,4).

Facilitates Chromatin Transcription (FACT) is a histone chaperone that facilitates DNA replication, transcription, and heterochromatin formation (5,6). FACT directly interacts with chromatin. Unlike other histone chaperones, FACT binds both DNA and histones in a nucleosome and subnucleosomal intermediates such as tetrasome and hexasome (5,7). Through these binding abilities, FACT preserves the chromatin structure during passage of DNA and RNA machinery through the gene locus (5,6). Therefore, FACT mutants exhibit a cryptic transcription phenotype, and FACT depletion in fission yeast results in defective heterochromatin (8–10).

FACT is a heterodimeric complex conserved from human to yeast. Yeast FACT is comprised of Spt16 and Pob3. Spt16 is composed of the N-terminal domain (NTD), dimerization domain (DD), middle domain (MD), and C-terminal domain (CTD). Pob3 is composed of an N-terminal and dimerization domain (N/DD), middle domain (MD), and C-terminal domain (CTD). Spt16 and Pob3 are associated through their dimerization domains. The cryo-EM structure of FACT bound to a hexasome showed that the DD complex is situated on the dyad axis of the nucleosome and two MDs interact with H3-H4 on both sides and two CTDs hold H2A-H2B via their acidic binding motif (11,12). In contrast, Spt16-NTD was not observed in the FACT-hexasome structure. The Spt16-NTD, which has a conserved peptidase fold (13,14), protrudes from the FACT-chromatin complex and its function in nucleosome reorganization remains unclear (11,15).

In the chromatin, FACT interplays with other proteins including RNA polymerase, chromatin remodelers, and histone chaperones (16–19). Abo1 has been identified as one of FACT’s interaction partners (20–26). Abo1 is one of the histone chaperones that bind histone H3-H4 and regulates the chromatin dynamics (20,27).

Abo1 is well-conserved from human to yeast. Fission yeast Abo1 enhances chromatin density and is required for heterochromatin formation (25,28). Interestingly, human Abo1, ATAD2 and budding yeast Abo1 homolog Yta7 also regulate nucleosome density but differ in their effect on chromatin from fission yeast Abo1. Abo1 belongs to the AAA+ family and forms a hexameric structure. Abo1 consists of an N-terminal region (NTR), AAA+ domains 1 and 2 (AAA1, AAA2), Bromodomain (BRD), and a C-terminal domain (CTD). The NTR is a long loop with the highest sequence variability among its homologues. Unlike other homologues, which commonly feature an acidic stretch within their NTR (29,30), Abo1 has a shorter segment rich in acidic residues between about 40-160 amino acids. AAA1 and AAA2, the most conserved regions, form a central two-tiered ring structure. BRD is a conserved protein fold that recognizes acetylated lysine. Our previous works showed that the Abo1 hexameric ring structure undergoes a conformational change upon ATP hydrolysis. The central pore of Abo1 hexamer holds the histone tail and loads H3-H4 onto DNA in an ATP-dependent manner (26).

Previous works show the genetical relationship between Abo1 and FACT. In fission yeast, depleting Abo1 derepresses cryptic transcripts that appeared also in *spt16* mutant cells (25). In addition, human Abo1, ATAD2 knock-out led to the stalling of FACT at the transcription starting site (TSS) (24). FACT subunit mutants alter the chromatin distribution profile of Abo1 suggesting that FACT works with Abo1 to regulate chromatin structure at genes (20,25). However, it is unclear whether the histone-loading activity of Abo1 cooperates with FACT or alternatively, Abo1 may have another function related to FACT.

In this study, we dissect the interplay between FACT and Abo1 via biochemical and single-molecule fluorescence imaging assay. Our data reveal that FACT and Abo1 directly interact with each other and that Abo1 facilitates the dissociation of FACT from tetrasome and nucleosome, implying that Abo1 may be involved in FACT recycling. Our findings provide insight how the interplay between histone chaperones regulate the chromatin dynamics.

## Material and Methods

### Construct cloning

*Schizosaccharomyces pombe (S. pombe)* Abo1 (1-1190 a.a.) and Abo1ΔN (243-1190 a.a.) were cloned into a modified pFastBac1 vector that includes TEV protease cleavage site and 1× flag tag at C-terminus. Abo1 pore mutant W345A and E385A were generated with KOD plus mutagenesis kit (Toyobo) based on the pFastBac1-Abo1-flag construct. The full-length of *S. pombe* Spt16 (1-1019 a.a.) and Pob3 (1-512 a.a.) were codon-optimized for Spodoptera frugiperda 9 (Sf9) insect cells and respectively cloned into a vector pFastBacTM HT B, which contains N-terminal 6×His tag followed by TEV protease cleavage site. *S.pombe* Spt16ΔC (1-937 a.a.) and Pob3ΔC (1-446 a.a.) were generated with KOD plus mutagenesis kit (Toyobo) based on the full-length constructs. To co-express Spt16 and Pob3 in Sf9 cells, the Pob3 or Pob3ΔC were cloned into pFastBac1 with no tag. For the single molecule imaging assay, *S. pombe* Abo1, Spt16, and Spt16ΔC were cloned into a modified pFastBac1 vector that included N-terminal 3× flag tag. For the GST pull-down assay, Spt16 domains were cloned into vector pGEX-4T1 containing an N-terminal GST tag. Spt16-NTD mutants K264A, R364A/F365A, and K264A/R364A/F365A were generated with KOD plus mutagenesis kit (Toyobo) based on the pGEX-4T1-Spt16-NTD construct.

### Protein purification

Full-length Abo1, Abo1ΔN, Abo1_W345A_, and Abo1_E385A_ were expressed in insect cells. After harvest, cell pellets were resuspended in lysis buffer (50 mM Tris-HCl pH 8.0, 300 mM NaCl, and 5 % (v/v) glycerol), with cOmpleteTM protease inhibitor cocktail (Roche). Cells were lysed by 4 cycles of freezing and thawing. The cell lysate was centrifugated at 15,000 rpm (∼ 32,500 xg) for 2 h. The supernatant containing Abo1 was incubated with anti-FLAG M2 resin (Sigma) at 4 ℃ for 2 h with rotating. The protein-bound beads were collected into a column and washed with the lysis buffer. The protein was eluted with elution buffer (lysis buffer supplemented with 0.4 mg/ml flag peptide (Apeptide). The eluted protein was diluted to 100 mM NaCl with 50 mM Tris-HCl buffer and applied to a HiTrap 5ml Q column (Cytiva). The protein was eluted using gradient elution from 50 mM NaCl to 1 M NaCl over 20 CV. Abo1 fractions were re-purified on Superose 6 increase 10/300 GL (Cytiva) column, pre-equilibrated in final buffer (20 mM HEPES pH 7.5, 200 mM NaCl, and 5 % (v/v) Glycerol).

*S. pombe* FACT was generated by co-expressing 6×His-tag-Spt16 and Pob3 in insect cells; the C-terminal deleted FACT (FACTΔC) by co-expressing 6×His-tag-Spt16_Δ938-1019_ and Pob3_Δ447-512_; After harvest, cell pellets were resuspended in the lysis buffer with a protease inhibitor cocktail (Roche). Cells were lysed by 4 cycles of freezing and thawing and the cell lysate was centrifugated at 15,000 rpm (∼ 32,500 xg) for 2 hours. The 6×His-tag-FACT complex was captured with Ni-NTA beads (Qiagen) while rotating at 4 ℃ for 1 hour. Following this, the beads were washed with wash buffers (50 mM Tris-HCl pH 8.0, 5% (v/v) Glycerol, and 20 mM imidazole) with varying salt concentrations of 300, 500, and 1000 mM NaCl as each 3CV. After equilibrating with wash buffer containing 300 mM NaCl, the bound proteins were eluted using elution buffer (lysis buffer supplemented with 100 mM imidazole). Fractions containing FACT were pooled and the His-tag was removed by TEV protease during overnight in dialysis buffer (lysis buffer supplemented with 1 mM DTT and 0.5 mM EDTA). The cleaved His-tag and TEV protease were eliminated by recapturing with Ni-NTA beads. The FACT proteins were diluted to 100 mM NaCl with 50 mM Tris-HCl buffer and applied to a HiTrap 5 ml Q column (Cytiva) and eluted using gradient elution from 50 mM to 1 M NaCl over 20 CV. The protein was further purified with Superdex 200 26 600 GL column (Cytiva) equilibrated with the final buffer (20 mM HEPES pH 7.5, 200 mM NaCl, and 5 % (v/v) Glycerol). Spt16 or Pob3 alone were respectively expressed 6×His-tag-Spt16 or 6×His-tag-Pob3 and purified as same with FACT complex purification.

For single molecule imaging assay, 3×FLAG-Spt16 was co-expressed with Pob3 in insect cells and the 3×FLAG-FACT complex was purified in the same way as FLAG-tagged Abo1.

### Native PAGE analysis

For Abo1 binding to FACT complexes and the FACT subunits, 100 nM of Abo1 was titrated into the increasing amount of FACT complexes and its subunits in a binding buffer (20 mM HEPES pH 7.5, 150 mM NaCl, and 2.5% glycerol) for 20 min at room temperature (RT, 23 ℃). The binding was analyzed with 3-12% Bis-Tris Native PAGE (Invitrogen) followed by Coomassie blue staining.

### GST pull-down assay

GST-tagged Spt16 domains were expressed in E. coli BL21 (DE3) RILP strain at 18 ℃ in the presence of 0.5 mM IPTG for 18 hours, and harvested in the lysis buffer supplemented with 1 mM PMSF. Cells were lysed by sonication and cleared by centrifugation at 15,000 rpm (∼ 21,000 xg) at 4 ℃ for 10 min. The supernatant was incubated with GST beads for 1 hour and washed by 3 CV of wash buffers (50 mM Tris-HCl pH 8.0, and 5% (v/v) Glycerol) with varying salt concentrations of 300, 500, and 300 mM NaCl. The beads bound with the proteins were equilibrated with the binding buffer (20 mM HEPES pH 7.5, 150 mM NaCl, and 2.5% glycerol) and incubated with 0.4 μM of Abo1 at 4 ℃ for 30 min with rotating. After removing flow-through, the beads were washed 5 times by 10 CV volumes of wash buffer containing 300 mM NaCl. Resin-bound proteins were resolved by 4-15% gradient SDS PAGE. All other GST-pulldown experiments were done with in a similar way.

### Fluorescence labelling of histone proteins

All recombinant *X. laevis* histones, including H2A, H2A_T120C_, H2B, H3, H4, and H4_T71C_, were purified using published protocol [50, 51]. To make fluorescence labeled H3-H4, dissolved H4_T71C_ in an unfolding buffer (20 mM Tris pH 7.5, 6 M Guanidine hydrochloride, 1 mM EDTA, and 0.5 mM TCEP) was combined with a 15-fold molar excess of Cy5 maleimide dye and incubated at RT for 5 hours. Excess dye was removed by dialysis, and the protein was refolded with wild-type H3 in a refolding buffer (20 mM Tris pH 7.5, 2 M NaCl, 1 mM EDTA, and 5 mM 2-mercaptoethanol). The refolded H3-_Cy5_H4_T71C_ was purified using a Superdex200 increase column (Cytiva) equilibrated with the refolding buffer. To make fluorescence labeled H2A-H2B, H2A_T120C_ was labeled with Alexa488 or Cy5 and refolded with wild-type H2B.

### In vitro tetrasome reconstitution

Lyophilized H3 and H4 were dissolved in unfolding buffer (20 mM Tris-HCl pH 7.5, 6 M Gu-HCl, and 5 mM DTT). H3 and H4 were mixed in equimolar concentration and dialyzed with refolding buffer (10 mM Tris-HCl pH 7.5, 2 M NaCl, 1 mM Na-EDTA, and 5 mM 2-Mercaptoethanol) at 4℃ overnight. Histone H3-H4 tetramer was applied to Superdex 200 26 600 GL column (Cytiva) pre-equilibrated in the refolding buffer. The peak fractions were concentrated using Amicon Millipore centrifugal concentrator, and stored at 4℃.

Tetrasome DNA was purified following a previously published protocol (31). The DNA sequence (−40 to 33) of the 601-positioning sequence was cloned into the pUC19 vector with the EcoRV enzyme site, and the DH5α strain was transformed with the resulting plasmid. The transformed cells were cultured in TB media, and the amplified plasmid was purified using Sol I, Sol II, and Sol III. The remaining RNA was treated with RNase A at 37℃ overnight and processed through a Sepharose 6 column (Cytiva). After purification, the 8 copies of DNA were cut using EcoRV (Enzynomics) at 37℃ overnight. To stop the enzymatic reaction and isolate DNAs, phenol extraction and isopropanol precipitation were performed. The 80bp DNA was cleared from the vector through size exclusion chromatography using a Sephacryl 1000 column (Cytiva). Tetrasome was reconstituted in vitro according to the published protocol [52]. A mixture of purified 80bp DNA and H3-H4 tetramer was prepared at a ratio of 1.0 to 1.3, depending on the batch of DNA or tetramer used. The salt concentration in the mixture was gradually reduced from 2M to 150 mM NaCl over 16 h using a dual peristaltic pump. The resulting mixture was subsequently dialyzed in tetrasome buffer (10 mM Tris-HCl pH 7.5, 150 mM NaCl, 1 mM EDTA, and 1 mM DTT) for 16 hours and tetrasome assembly was confirmed via 5% TBE native PAGE analysis.

### Electrophoretic mobility shift assay (EMSA)

In conducting the FACT dissociation assay with TBE native PAGE, FACT and H2A-H2B were mixed at a 1:2 molar ratio and incubated for 30 min at RT. Subsequently, 200 nM of the FACT/H2A-H2B complex was added to a reaction tube containing 100 nM of tetrasome in assay buffer (10 mM Tris-HCl pH 7.5, 150 mM NaCl, and 1 mM DTT) for 30 min at RT. 200 nM of Abo1 was then supplemented to each reaction tube with 1 mM of ATP at RT and incubated for 30, 60, 90 min. For negative control, 200 nM of Abo1 was added to the reaction tube with 4 mM EDTA at RT for maximal time condition, 90 min. After the reaction, 50% sucrose was added in 5× concentration as a loading buffer and the sample was loaded onto 0.2×TBE 5% acrylamide gel. Before running, the TBE gel was pre-run at 150 V for 60 min, and run at 120 V for 60 min. Cy5 fluorescence signal was detected at 695 nm for the detection of histone H2A or H4, followed by DNA detection staining with SYBR Gold (Thermo Fisher Scientific).

### Tetrasome formation assay

50 nM of wild-type or tailless H3-H4 dimer were pre-incubated with CAF-1 at different concentrations (0, 100, and 200 nM) in CAF-1 buffer (25 mM Tris-HCl pH 7.5, 150 mM NaCl, 1 mM EDTA, 0.02% Tween 20, and 0.5 mM TCEP) at 23°C for 10 min. 50 nM of 147 bp Widom 601 DNA was incubated with the CAF-1-H3-H4 at 23°C for 30 min for tetrasome formation. The samples were analyzed in 6% non-denaturing PAGE with SybrGOLD (S11494, Invitrogen) staining. The fluorescence was imaged by Typhoon RGB (Cytiva).

### AlphaFold prediction

The primary amino acid sequence of *S. Pombe* Abo1 (UniProt ID: O14114) and Spt16 (UniProt ID: O94267) were obtained from the UniProt database. The N-terminal region of Abo1 (1-240 a.a.) and Spt16-NTD domain (1-442 a.a.) were submitted to the AlphaFold v2.0 model using default parameters. The resulting five structural models were obtained and visualized using PyMOL software. RMSD values were calculated to assess the structural similarity among the models. When comparing only the Spt16-NTD and FIN helix regions, the RMSD values between models range from 0.209 to 0.664.

### Single-molecule dissociation assays

#### DNA preparation for dissociation assay

All oligomers were synthesized by Bioneer (South Korea) and listed in Table S1. For biotinylated 80 bp Widom 601 DNA, complementary oligomers (Widom601_80_biotin and Widom601_80_comp) were hybridized in Lipid buffer (20 mM Tris-HCl pH 8.0, and 100 mM NaCl). Biotinylated 147 bp Widom 601 DNA was prepared by PCR amplification with primers (Widom601_147_F_biotin and Widom601_147_R) (Table S1). *Total internal reflection fluorescence microscope (TIRFM):* Prism-type total internal reflection fluorescence microscope (TIRFM) was custom-built with Nikon Eclipse Ti-2. Solid-state 488-nm and 637-nm lasers (200 mW, OBIS, Coherent Laser) were used to excite quantum dot (QD) and Cy5, respectively. FLAG-antibody-conjugated QD (605 nm emission), which was prepared using a commercial kit (S10469, Invitrogen). Laser beam was incident through a prism to increase the incident angle beyond the critical angle, so that the laser beam was totally reflected at the interface between the surface of fused-silica slide and buffer to generate evanescent field with approximately 250 nm skin depth. The fluorescence signal from fluorophores was collected by a 60× water-immersion objective lens (CFI Plan Apo VC 60XWI; Nikon) and imaged by an EM-CCD camera (iXon 897; Andor Technology). To block the laser beam, a long-pass filter suitable for each fluorophore was used. The imaging data were collected by the NIS-Element software (Nikon).

#### Lipid preparation

Liposome was prepared as previously described (32,33). 10 mg/ml DOPC (1,2-dioleoyl-sn-glycero-phosphocholine) (850375P-500 MG, Avanti), 0.8 mg/ml mPEG 2000-DOPE (1,2-dioleoyl-sn-glycero-3-phosphoethanolamine-N-[methoxy(polyethylene glycol)-2000]) (880130P-25MG, Avanti), and 0.5% biotinylated-DOPE (1,2-dioleoyl-sn-glycero-3-phosphoethanolamine-N-(cap biotinyl) (sodium salt), 870273, Avanti) were dissolved in chloroform and mixed in a glass vial. For single-molecule imaging of FACT or tetrasome dissociation, we used an unbiotinylated lipid bilayer by omitting biotinylated-DOPE. The lipid was dried on the side of the glass bottle wall by careful evaporation of chloroform with nitrogen gas. The glass vial was placed under vacuum overnight to completely remove residual chloroform. Lipid buffer was then added to the dried lipid mixture for lipid hydration for at least 2 hours. The hydrated lipid was vortexed for 2 ∼ 3 min to form large multilamellar vesicles, which were converted into small unilamellar vesicles using sonication (amplitude: 50%, pulse on: 90 sec, pulse off: 10 min, total running time: 4 min 30 sec, Qsonica). After sonication, the liposome solution was filtered using a 0.22 μm nylon syringe filter to remove the debris from a sonication tip.

#### Flowcell preparation for single-tethered DNA curtain assay

Nano-trenches were fabricated on fused-silica slides followed by the previous protocol (34). Briefly, fused-silica slides with two holes were cleaned in piranha solution (3:1 mixture of concentrated 96% H_2_SO_4_ with 30% H_2_O_2_). On the surface of the slides, aluminum (Al) was evaporated and 950K PMMA for e-beam resistor was spin-coated. Nano-trench patterns were drawn on the PMMA layer by electron-beam lithography and then developed in a developing solution (1:3 MiBK:IPA). The slide surface coated with PMMA and Al layers was etched to form nano-trenches by inductively coupled plasma (ICP)-RIE using a mixture of Cl_2_ and BCl_2_ gases. The remaining Al and PMMA layers were removed by Al etchant (AZ-MIF300) and piranha solution (3:1 H_2_SO_4_: H_2_O_2_), respectively. The slides were cleaned in 2% Hellmanex III (Sigma) for at least one day, acetone for 30 min, and 1 M sodium hydroxide for 20 min, successively. The slides were rinsed with deionized (D.I.) water between each step. A microchannel was formed by gluing the patterned slide and a coverslip with a double-sided tape. Nanoports (IDEX) were attached around the holes to link the flowcell with a syringe pump-based fluidic system. The flowcell was rinsed with D.I. water and then with Lipid buffer. The lipid bilayer was built up on the microchannel surface by washing out free liposomes followed by 20 min incubation. BSA buffer (40 mM Tris-HCl pH 8.0, 50 mM NaCl, 2 mM MgCl_2_, and 0.4% BSA) was then added for further surface passivation. The lambda DNA (λ-DNA) tagged with biotin at one end was anchored on the biotinylated lipid via streptavidin. The flowcell was connected with the syringe-based fluidic system and then placed on TIRFM. Under continuous buffer flow, DNA molecules were moving along the flow and then aligned at the nano-trenches.

#### DNA curtain assays for tetrasome or nucleosome assembly

All DNA curtain experiments were performed at 23°C. The formation of DNA curtains was ensured in BSA buffer supplemented with 0.1× Gloxy and 1.6% glucose by staining λ-DNA with an intercalating dye YOYO-1, which was excited by the 488 nm laser (35). After YOYO-1 was removed from λ-DNA by washing with 20 mM MgCl_2_ and 300 mM NaCl using an injection system. Then, to test tetrasome assembly, 1.5 nM Abo1 (or CAF-1) and 1.5 nM FACT (or FACTΔC) were pre-incubated in Reaction buffer (50 mM Tris-HCl pH 8.0, 100 mM NaCl, 2 mM DTT, 2 mM MgCl_2_, 0.1× Gloxy, and 1.6% glucose) supplemented with 1 mM ATP on ice for 15 min. 6 nM Cy5-labeled H3-H4 dimers (H3-_Cy5_H4) were then added to Abo1 (or CAF-1) and FACT (or FACTΔC) mixture and incubated on ice for 15 min. The protein mixture was injected into the flowcell and incubated for 15 min. The binding of H3-H4 dimers to λ-DNA was visualized by a 637 nm laser. For nucleosome assembly, 2 nM Abo1 (or CAF-1) and 5 nM H3-_Cy5_H4 were pre-incubated in Reaction buffer I supplemented with 1 mM ATP on ice for 15 min. The protein mixture was injected into the flowcell and incubated for 15 min. After H3-H4 dimers bound to λ-DNA were imaged by a 637 nm laser, 2 nM FACT and 5 nM Alexa488-H2A-H2B, which were pre-incubated on ice for 15 min, were added to the flowcell. The assembly of H2A-H2B into tetrasome was visualized under the illumination of a 488 nm laser. Except imaging procedure, all lasers were turned off to prevent the photobleaching of Cy5 or Alexa488.

#### Analysis of single-molecule DNA curtain assay

All DNA curtain images were taken by NIS-Elements (Nikon) and analyzed by ImageJ (NIH) in TIFF format. To count the number of H3-H4 dimers bound to DNA, the fluorescence intensity of individual Cy5 puncta was divided by single Cy5 fluorescence intensity (minimum intensity of Cy5 punctum).

#### Single-molecule fluorescence imaging for FACT or tetrasome dissociation

A flowcell with microchannel was assembled by gluing a fused-silica slide and a coverslip using double-sided tape. 0.2 mg/ml streptavidin (S888, Invitrogen) in Lipid buffer was nonspecifically adsorbed on the flowcell surface followed by 10 min incubation. The liposome without biotin was injected into the flowcell with 10 min incubation. The lipid bilayer was promoted on the slide surface for 20 min after washing out excessive liposomes by Lipid buffer (32). The lipid bilayer passivated the surface to prevent nonspecific binding of proteins. 1 nM of biotinylated 80 bp or 147 bp Widom 601 DNA was anchored on the surface via streptavidin. Unbound DNA was washed out by Lipid buffer. 1 nM *S. cerevisiae* CAF-1 and 1 nM *X. laevis* H3-H4 dimers were pre-incubated on ice for 15 min in Reaction buffer supplemented with 1 mM ATP if needed. The mixture was injected into the flowcell and incubated at 23°C for 15 min to form a tetrasome. For the FACT dissociation experiments, *S.pombe* FACT having 3×FLAG at N-terminus was fluorescently labeled by FLAG-antibody-conjugated Qdot (605 nm emission), which was prepared with a commercial kit (S10469, Invitrogen). FACT and FLAG-antibody-Qdot were mixed at 1: 1 molar ratio and incubated on ice for at least 10 min. In the following procedure, residual proteins were washed out in Reaction buffer II before the next proteins were injected. Then 30 pM of FACT-Qdot was added into the flowcell and incubated at 23°C for 30 min. Finally, 0.5 nM of full-length Abo1 (FLAbo1) or FLAbo1 mutant was added into the flowcell. After 30 min incubation, fluorescence images were taken under TIRFM with 488 nm laser. To test the tetrasome destabilization by FACT or FLAbo1, we followed the same procedures as above except that FACT was not tagged by Qdot but H3-H4 dimer was labeled with Cy5 at H4 (T71C) (26). For the tetrasome formation, 300 pM of CAF-1 and 750 pM of H3-_Cy5_H4 were pre-incubated and then injected into the flowcell with Widom 601 DNA. Then 5 nM unlabeled FACT and 5 nM FLAbo1 were added successively. Cy5 fluorescence was imaged under TIRFM with 637 nm laser.

#### Analysis of single-molecule dissociation assay

All single-molecule fluorescence images were taken by NIS-Elements software (Nikon) and then transformed into TIFF format. All image data were analyzed using ImageJ (NIH). The first 5 frames of each image were averaged. Fluorescence puncta were counted using an ImageJ plug-in “Find maxima.”

### Fission yeast culture and genetic manipulations

Standard culturing techniques were used in this study. Tetrad dissections and standard genetic crosses were used to obtain *S. pombe* strains. PCR-based techniques were used to construct gene deletion and epitope-tagged strains. Abo1 protein missing first 242 amino acids *(abo1ΔN*) was expressed from its native locus. N-terminal truncation was achieved by homologous recombination with chemically synthetized construct lacking the deleted section of *abo1* gene and assisted by CRISPR-Cas9 cleavage that targeted sequence inside the deleted regions.

### Dilution assay

Expression of *mat2::ura4^+^* reporter sensitized by deletion of proximal *REII* silencer was monitored by spotting serial dilutions on minimal PMG (*pombe* minimal medium) depleted for uracil or supplemented with *ura4^+^* counter-selective 5-fluoroorotic acid. Expression of *mat2* locus was also monitored by exposing cells grown on PMG to iodine vapors that stains brown the starch-like substance accumulated in spores.

### Chromatin Immunoprecipitation

Histone H3 lysine 9 methylation and Swi6/HP1 localization was analyzed in mid-log phase cells grown at 32°C in rich (YEA) media. Cells were treated with 3% paraformaldehyde (Sigma-Aldrich) and fixation quenched after 30 min by adding 125 mM glycine. Twice PBS-washed cells were re-suspended in lysis buffer (50 mM HEPES/KOH, pH 7.5, 140 mM NaCl, 1 mM EDTA, 1% Triton X-100, 0.1% sodium deoxycholate) supplemented with 1mM PMSF and Complete protease inhibitors (Roche), and immediately lysed by bead-beating (Biospec Mini-Beadbeater-16). Chromatin was sheared at 4°C to 0.4-0.6 kb by sonification (Bioruptor^+^, Diagenode) for 12 cycles 30 sec ON 30 sec OFF on low power settings, and after lysate pre-clearing by centrifugation at 1,500g (8 min., 4°C), 3% of cells extract was set apart as an input control sample. After lysate incubation with Sepharose 6B (Sigma-Aldrich), 2 μgs of H3K9me3 antibody (Ab 8898, Abcam) or in-house prepared Swi6 antibody was incubated overnight with pre-cleared lysate and immunoprecipitated chromatin recovered by incubation with 30 μls of 50% protein A slurry for 4 hours at 4°C. The slurry was washed twice with each: lysis buffer, lysis buffer supplemented with 0.5 M NaCl, wash buffer (10 mM Tris/HCl pH 8, 250 mM LiCl, 0.5% sodium deoxycholate, 0.5% IGEPAL and 1mM EDTA). After final wash with TE buffer, immunoprecipitated chromatin was eluted by repeated 25 min incubation at 65°C in 1×TE buffer supplemented with 1% SDS. The crosslink of immunoprecipitated sample and input control was reversed at 65°C overnight. DNA was column purified (Mini-elute QIAquick PCR purification kit, Qiagen) after sequential RNase A (4μg,) and proteinase K (40μgs) 30 min treatments at 37°C.

### ChIP qPCR

DNA recovery of targeted heterochromatic locus in ChIP and input DNA was quantified by qPCR analysis (iTaq Universal SYBR Green Supermix, BioRad) on QuantStudio^TM^ 3 real-time PCR machine and enrichment calculated using delta-delta Ct normalization to reference *leu1^+^* locus.

## Results

### Abo1 directly interacts with FACT through the N-terminal domain (NTD) of Spt16

Abo1 was identified as one of FACT interacting partners in *S. Pombe* (25). However, it was not clear whether Abo1 directly interacts with FACT. Therefore, we used native gel analysis to test the interaction between Abo1 and FACT. For this purpose, we generated recombinant Abo1 and FACT proteins (Figure S1A). Abo1 was incubated with FACT and the complex formation was analyzed by native PAGE. Unexpectedly, there was no significant change in gel mobility even in the excess amount of FACT (Figure 1A, lane 2-4), indicating that full-length FACT does not interact with Abo1 in our condition. Previously, FACT was shown to undergo conformational changes from compact to open forms when its C-terminal regions is altered (11,36). This C-terminal region seems to have an autoinhibitory activity that maintains the compact form of the whole FACT. When the C-terminus is deleted or bound by other proteins, the conformation of the whole FACT complex changes to an open form (36,37). Based on these observations, we hypothesized that the Abo1 binding region of FACT may be blocked in the compact conformation. Therefore, we attempted to investigate whether the open conformation of FACT can bind to Abo1. To make the open form of FACT, we generated FACTΔC (Spt16_ΔC938-1019_ and Pob3_ΔC447-512_) where the C-terminal regions of both FACT subunits were deleted. We then tested the binding between Abo1 and FACTΔC using native PAGE analysis. We incubated Abo1 with the increasing amount of FACTΔC and analyzed the complex formation by native PAGE (Figure 1A, lane 5-7). Interestingly, Abo1 form a complex with FACTΔC, suggesting that FACT in an open conformation directly interacts with Abo1.

**Figure 1.**
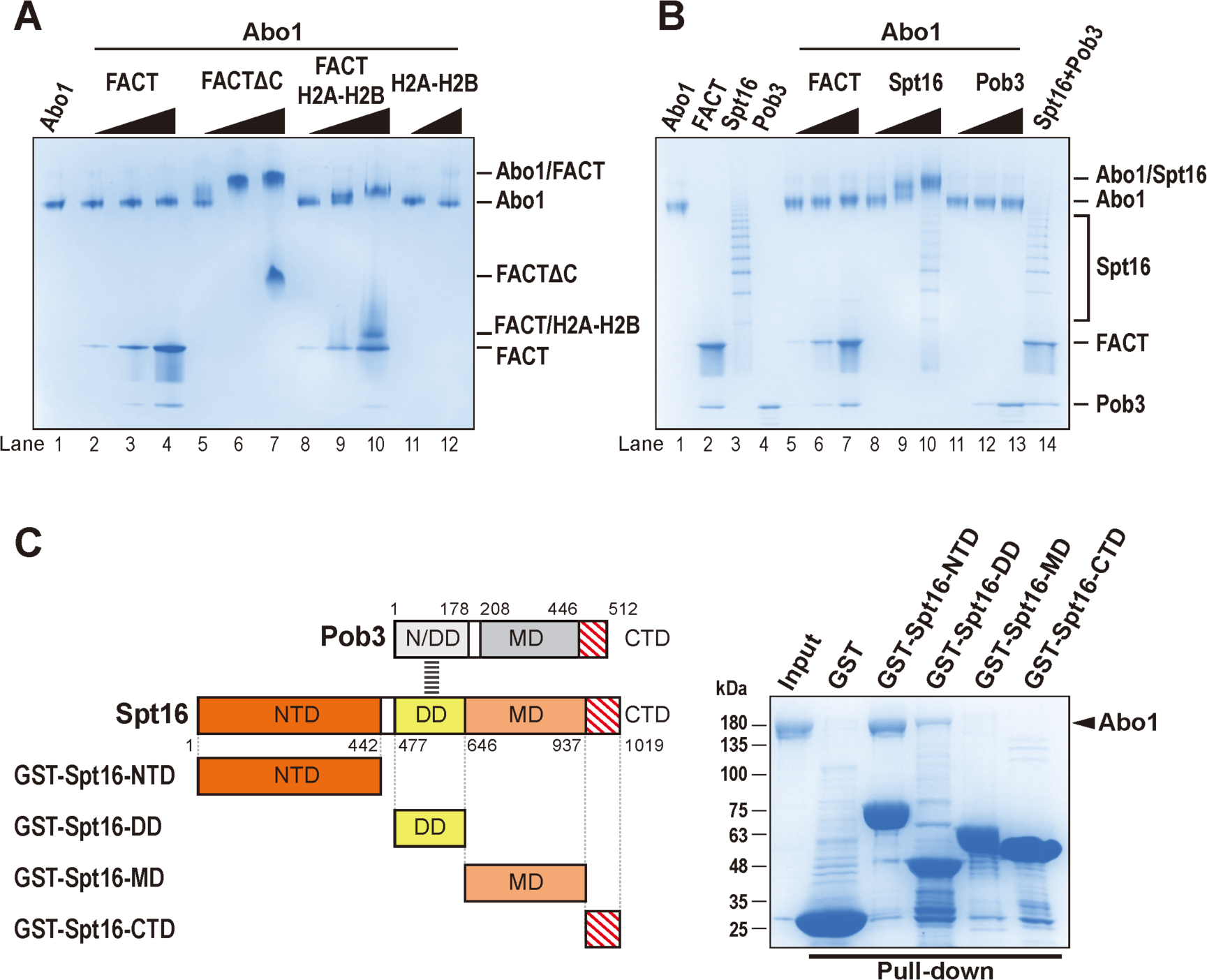
Abo1 directly binds to FACT via Spt16-NTD. (**A**) FACT does not interact with full-length FACT. FACTΔC consists of Spt16_Δ938-1019_ and Pob3_Δ447-512_. For FACT/H2A-H2B complex, H2A-H2B was pre-incubated with FACT at 1:2 molar ratio for 30 min at room temperature (RT). 100 nM Abo1 was incubated with the increasing amount (100, 500, or 1,500 nM) of FACT (lane 2, 3, 4), FACTΔC (lane 5, 6, 7), FACT/H2A-H2B (lane 8, 9, 10) or H2A-H2B (100, or 500 nM, lane 11, 12) for 1 hour at RT. The complex formation was analyzed using 3-12% Bis-Tris native PAGE and stained with Coomassie. (**B**) Spt16 subunit is responsible for Abo1 binding. 100 nM Abo1 was incubated with 100, 400, or 1,600 nM FACT (lane 5, 6, 7), Spt16 (lane 8, 9, 10), or Pob3 (lane 11, 12, 13) for 30 min at RT. Lane 14 shows the ability of FACT formation by mixing separately purified Spt16 and Pob3. The complex formation was analyzed using 3-12% Bis-Tris native PAGE and stained with Coomassie. (**C**) Spt16-NTD and DD bind to Abo1. (Left) A schematic diagram of *S. Pombe* FACT domains showing the GST-tagged domains used for the GST pull-down assay. (Right) GST pull-down assay with the GST-tagged domain for Abo1 binding. The bound proteins were analyzed by 4-15% SDS-PAGE followed by Coomassie staining.

The C-terminal region of FACT has highly acidic residues that interact with H2A-H2B (12). FACT conformation has been shown to be regulated by interacting with other proteins including histones under cellular conditions. Therefore, we asked whether H2A-H2B binding to the C-terminal regions of FACT also makes FACT capable of binding to Abo1. FACT was preincubated with H2A-H2B and Abo1 was incubated with the increasing amount of FACT/H2A-H2B complex. The complex formation was then analyzed by native PAGE (Figure 1A, lane 8-10), showing that in the presence of H2A-H2B, full-length FACT forms a complex with Abo1, These data suggest that FACT needs to undergo a conformational change to interact with Abo1.

FACT is a heterodimer complex of Spt16 and Pob3. We aimed to figure out which FACT subunit is responsible for Abo1 binding. To this end, Spt16 or Pob3 subunits were purified (Figure S1A) and the complex formation between each subunit and Abo1 was monitored by native-PAGE, showing that Spt16 subunit forms a complex with Abo1 but not Pob3 (Figure 1B, lane 8-13). To further narrow down the domain within Spt16 subunit, which is required for Abo1 binding, we generated a series of Spt16 domain in GST fusion forms (Figure 1C). We then examined the Abo1 binding of each Spt 16 domain by GST pull-down assay (Figure 1C). The GST pulldown assay showed that the N-terminal domain (NTD) of Spt16 strongly interacts with Abo1 while the dimerization domain (DD) marginally interacts with Abo1 (Figure 1D). Collectively, these data show that Abo1 and FACT directly interact via Spt16-NTD and Spt16-DD.

### Abo1 utilizes its N-terminal region (NTR) to interact with FACT

Next, we decided to dissect which region of Abo1 interacts with FACT. Abo1 belongs to the AAA+ family, and several AAA+ proteins utilize their N terminal domain for interaction with other proteins (38). Abo1 has a long N terminal region (NTR, 1-242 a.a.) containing many negatively charged amino acids (29) (Figure 2A). To investigate whether the Abo1 NTR is required for Spt16 binding, we generated a N-terminally truncated Abo1 (Abo1ΔN, 1-242 a.a. deletion). We then examined the binding of Abo1ΔN to FACT by GST-pulldown assay. The NTR deletion of Abo1 abrogated the ability to form a complex with FACTΔC, indicating the NTR of Abo1 is required for the interaction with FACT (Figure 2B). As Spt16-NTD and DD interact with Abo1 (Figure 1C), we tested whether Spt16-NTD or DD is responsible for interacting with the NTR of Abo1. We performed a GST pull-down assay using GST-Spt16-NTD or GST-Spt16-DD with Abo1ΔN, showing that GST-Spt16-NTD binds to full-length Abo1 but not Abo1ΔN while Spt16-DD shows binding to both full-length Abo1 and Abo1ΔN (Figure 2C). These data suggest that the NTD of Spt16 is mainly tethered to the NTR of Abo1.

**Figure 2.**
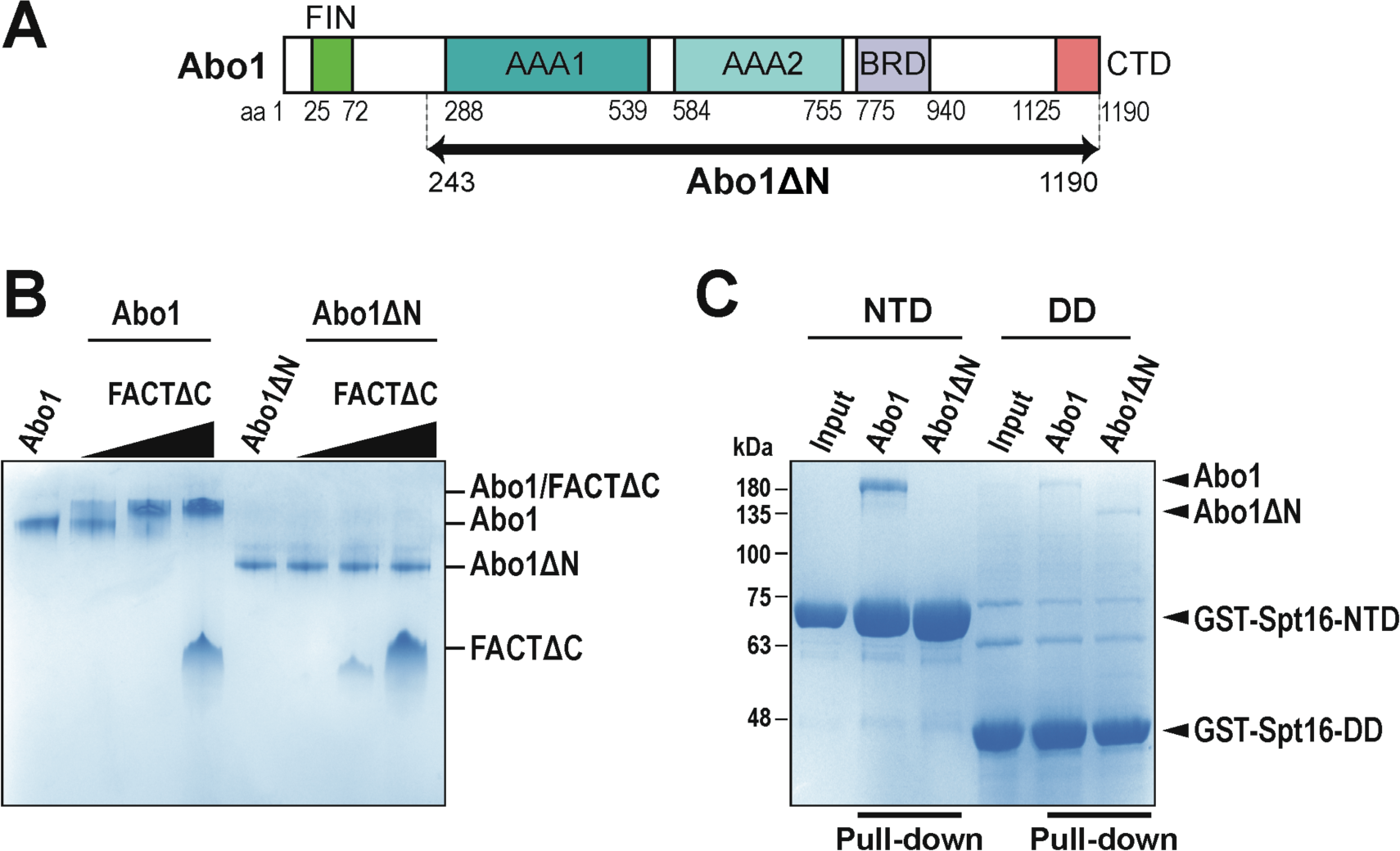
The N-terminal region of Abo1 is required for binding to the NTD in FACT. (**A**) A schematic diagram of Abo1 domains. Abo1 contains N-terminal region (NTR), AAA1, AAA2, bromodomain (Bromo), and C-terminal domain (CTD). FACT interacting (FIN) helix identified in this study is labeled. The black arrow line indicates the region of Abo1ΔN (243 to 1,190 a.a.). (**B**) The N-terminus of Abo1 is required for FACT binding. 100 nM Abo1 or Abo1ΔN was incubated with 100, 400, and 1,600 nM FACTΔC for 1 hour at RT and the binding was analyzed by 3-12% Bis-Tris native PAGE. (**C**) Spt16-NTD interacts with the Abo1 N-terminus. GST pull-down assay for Abo1 or Abo1ΔN. 0.5 μM of Abo1 or Abo1ΔN was incubated with beads containing GST-Spt16 domain. The proteins bound to GST-Spt16-NTD or DD were analyzed by 10% SDS PAGE followed by Coomassie staining.

Abo1 has a hexamer ring structure and undergoes a conformational change from an open to a closed ring upon ATP hydrolysis (26). To examine whether the conformational changes of Abo1 affects FACT binding, we compared the FACT binding with Abo1 in the presence of ATP or ADP and in the absence of nucleotide (Figure S2A). We observed that the interaction between Abo1 and FACT is not influenced by nucleotide status of Abo1. In addition, we also found that the binding between Abo1 and Spt16-NTD or DD was not affected by nucleotide binding status of Abo1 (Figure S2B) (Figure S2B). Taken together, these data demonstrate that Abo1 binds FACT via its N-terminus in an ATP-independent manner.

### FACT interacting (FIN) helix of Abo1 binds to the central cavity of Spt16-NTD

The structure of Spt16-NTD showed that is is composed of N- and C-lobes forming a central cavity between two lobes (13,14,39). The C-lobe adopts the aminopeptidase fold but Spt16-NTD does not have any enzyme activity (39). Instead, the central cavity is considered a platform for interacting with other proteins (39,40). Therefore, we reasoned that Spt16-NTD may utilize the central cavity to interact with the NTR of Abo1. To dissect the molecular interaction between Spt16-NTD and Abo1, we generated a predicted model of Spt16-NTD bound with a part of Abo1 NTR using AlphaFold (41) by providing the primary sequences of Spt16-NTD (1-442 a.a.) and NTR of Abo1 (1-242 a.a). Interestingly, the predicted model shows that a helix (25-72 a.a.) of Abo1 binds to the central cavity between the N- and C-lobes of Spt16-NTD (Figure 3A and S3A). Spt16-NTD includes a conserved basic patch in its central cavity (39), and the placement of the Abo1 helix in the model positions several acidic residues of the helix close to the basic patch of Spt16-NTD (Figure S3B). To validate the predicted interaction between the Abo1 helix and Spt16-NTD, we generated GST-Abo1 helix (25-72 a.a.) and performed GST pull-down with Spt16-NTD, showing that the Abo1 helix (25-72 a.a.) indeed interacts with Spt16-NTD (Figure 3B). We named the Abo1 helix (25-72 a.a.) as ‘FACT interacting (FIN) helix’. Based on the predicted model, we also mapped residues, which may be involved in the interactions between Spt16-NTD and FIN helix. K264 and R365 of Spt16-NTD are proximally located with E55 and E58 of the FIN helix at a distance of 2.7 Å, suggesting that they form electrostatic interactions. In addition, F364 of Spt16-NTD is positioned near F51 of Abo1, which may form a hydrophobic interaction (Figure 3A). To confirm the interaction interface between Spt16-NTD and Abo1 FIN helix, we generated mutants (K264E, F364A/R365E, or K264E/F364A/R365E) of Spt16-NTD (Figure 3B) that would disrupt the interaction. We then conducted a GST pull-down assay with GST-Abo1-FIN helix for Spt16-NTD and its mutants. We found that Spt16-NTD mutants showed substantially decreased binding (Figure 3B), suggesting that FACT utilizes the central cavity in Spt16-NTD to interact with the FIN helix in Abo1.

**Figure 3.**
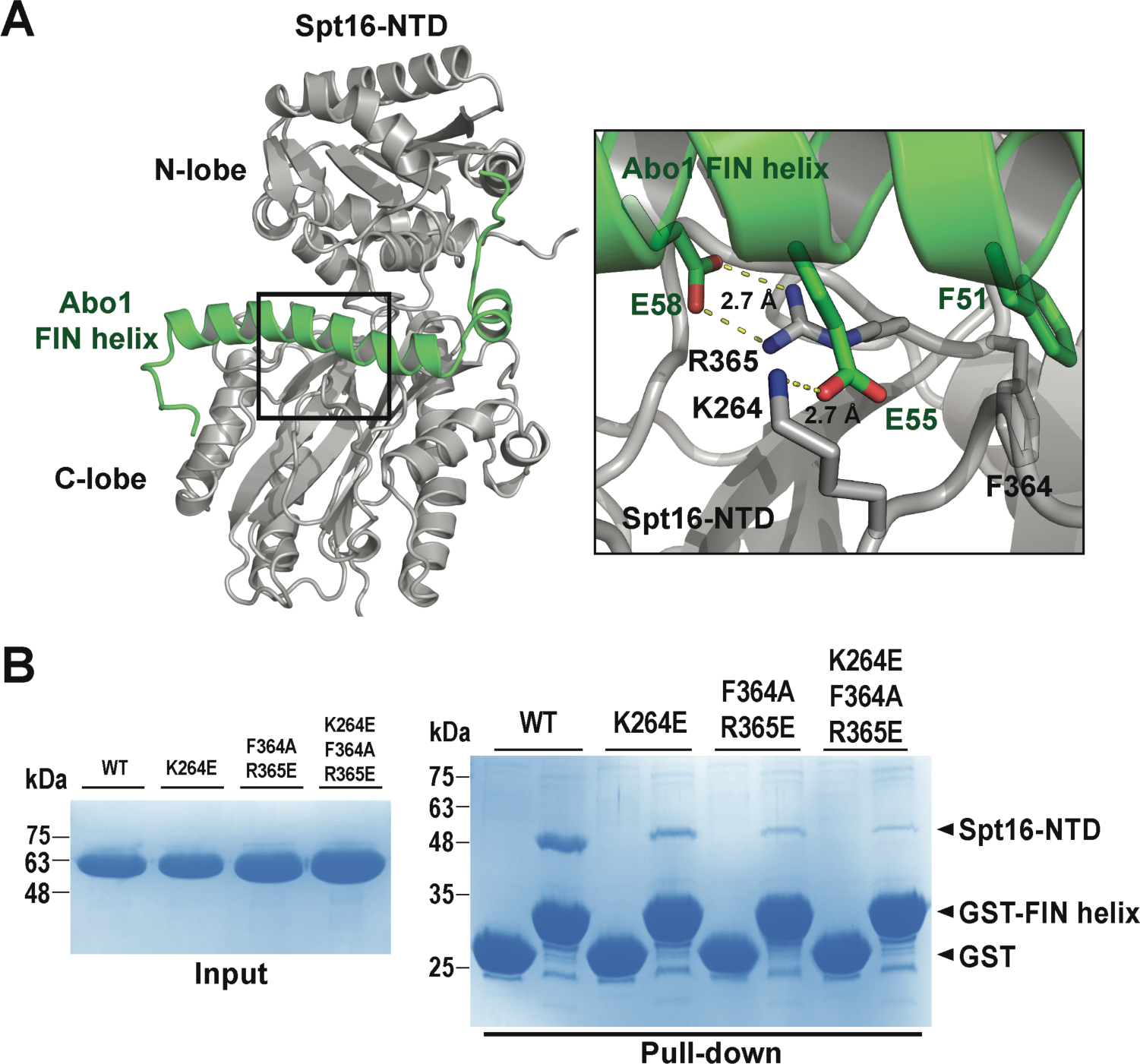
Structural analysis on the interaction between Spt16-NTD and the Abo1-FIN helix. (**A**) AlphaFold prediction model of Abo1-NTR and Spt16-NTD. (Left) Abo1 FIN helix (25-72 a.a.) was colored in green and Spt16-NTD (1-442 a.a.) was colored in gray. (Right) The detailed interaction between FIN helix and Spt16-NTD. The predicted model shows that K264 and R365 in Spt16-NTD electrostatically interact with E55 and E58 in FIN at a distance of 2.7 Å and that F264 of Spt16-NTD has hydrophobic interaction with F51 in FIN helix. (**B**) Mutation analysis of Spt16-NTD binding to the Abo1 FIN helix. (Left) Mutations in Spt16-NTD disrupt Abo1 binding. Purified Spt16-NTD proteins were visualized by SDS-PAGE (WT, K264E, F364A/R365E, K264E/F364A/R365E). (Right) GST pull-down assay with GST-Abo1-FIN helix and Spt16-NTD mutants. 0.5 μM of WT or the mutants of Spt16-NTD were incubated with GST-Abo1-FIN helix. The binding between Abo1 FIN-helix and Spt16-NTD mutants was analyzed by SDS-PAGE.

### Abo1 facilitates the dissociation of FACT from chromatin in an ATP-dependent manner

In our previous study, we showed that Abo1 loads histone H3-H4 on DNA in an ATP-dependent manner (26). We first hypothesized that FACT might modulate the histone-loading activity of Abo1. To test this hypothesis, we employed a DNA curtain assay, which allows direct observation of fluorescently labeled proteins using total reflection fluorescence microscopy (TIRFM). To visualize histones, we labeled histone H3-_Cy5_H4, where H4_T71C_ was labeled with Cy5 maleimide dye. We incubated Abo1 and H3-_Cy5_H4 and injected it into a microfluidic chamber where Lambda DNA molecules were aligned and stretched at a nano-barrier. Using this DNA curtain system, we compared the amount of H3-H4 loaded onto DNA by Abo1 in the presence or absence of FACT, indicating that the histone loading activity of Abo1 is not modulated by FACT (Figure S4A). In addition, we also showed that FACT does not modulate the histone H3-H4 loading activity of chromatin assembly factor 1 (CAF-1), a well-known H3-H4 histone chaperone responsible for H3-H4 tetrasome assembly (42–44).

We further investigated whether FACT may load H2A/H2B to assemble nucleosome following H3/H4 loading by Abo1. To visualize two types of histones, we labeled H2A_T120C_ with Alexa488 and H4_T71C_ with Cy5, then refolded them into _Alexa488_H2A/H2B and H3/_Cy5_H4, respectively. Pre-incubated CAF-1 and H3/_Cy5_H4, or Abo1 and H3/_Cy5_H4 were added to the microfluidic chamber to load H3/_Cy5_H4onto tethered DNA molecules. Loading of H3/_Cy5_H4 was confirmed by formation of fluorescent puncta (shown in red in Figure S4B). To examine nucleosome assembly, we injected FACT _Alexa488_H2A-H2B complex into the same microfluidic chamber and observed co-localization of Alexa488 and Cy5 fluorescence signals. The results show a significant amount of _Alexa488_H2A-H2B co-localized with H3/_Cy5_H4 loaded by CAF-1, but little with H3/_Cy5_H4 loaded by Abo1 (Figure S4B), concluding that Abo1 and FACT do not cooperate in nucleosome assembly.

A previous study has shown that depletion of human Abo1 leads to the arrest of FACT at the transcription start site (TSS), suggesting that Abo1 impacts FACT distribution profile of chromatin (24). Consequently, we sought to investigate whether Abo1 affects the loading or unloading of FACT from chromatin. To test this hypothesis, we designed a single-molecule dissociation assay (Figure 4A). First, we generated the recombinant FACT which has 3× FLAG tag at the N-terminus of Spt16 subunit. The 3×FLAG-FACT was then fluorescence-labeled with anti-flag-QD and pre-incubated with H2A-H2B. As FACT dynamically assembles and disassembles the nucleosome, we imposed constraints on DNA length and attempted to mimic the ‘chromatin-bound FACT’, an intermediate structure in which FACT sits on a tetrasome holding H2A-H2B by its C-terminal tail (5,11). We immobilized short 80 bp DNA on the surface of a flowcell and assembled H3/H4 tetrasome by CAF-1, followed by addition of pre-assembled QD-FACT/H2A-H2B to induce the formation of the QD-FACT/H2A-H2B/tetrasome complex.

**Figure 4.**
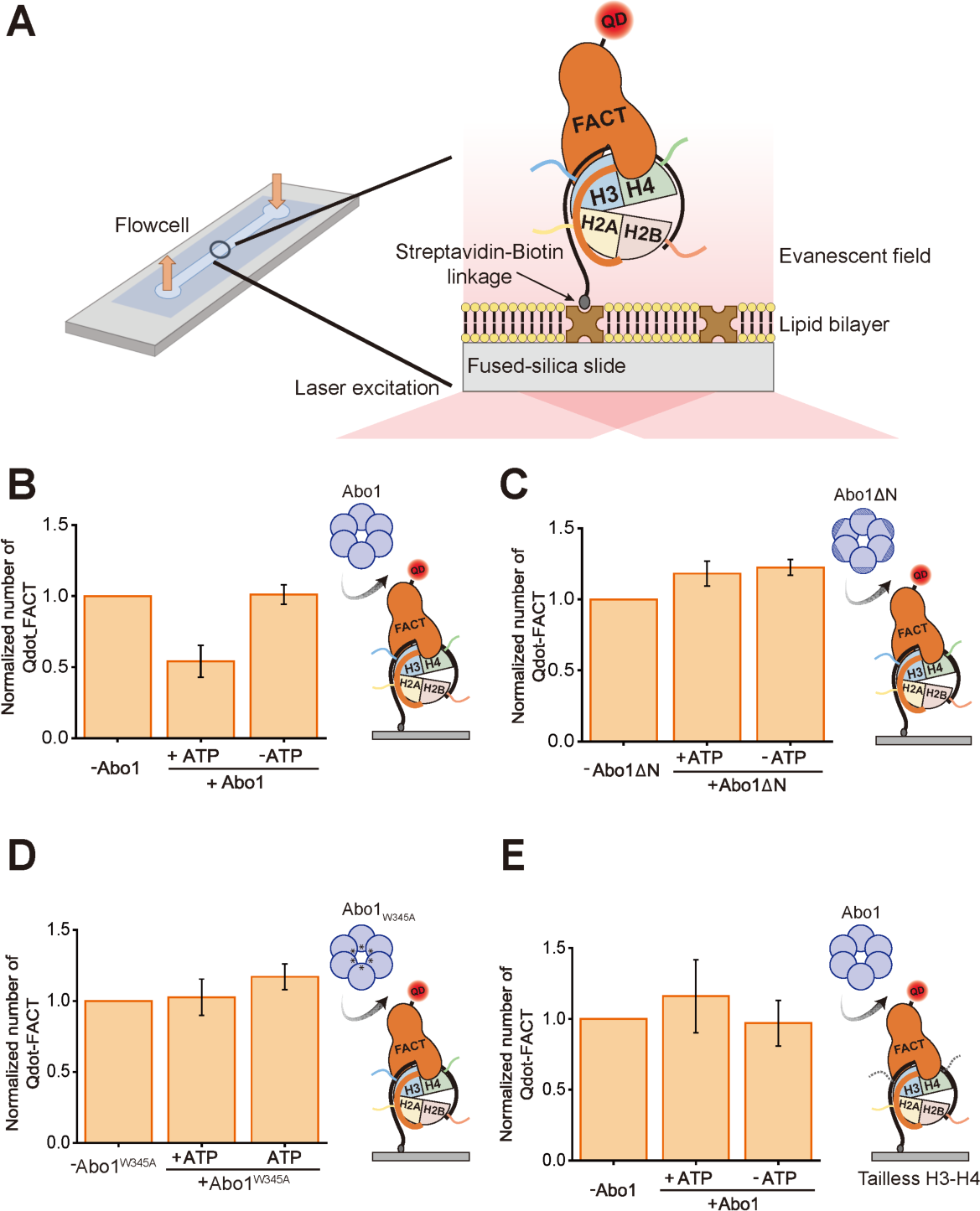
Abo1 facilitates the dissociation of FACT from tetrasome in an ATP-dependent manner. (**A**) A scheme of single-molecule fluorescence imaging of FACT dissociation by Abo1 using total internal reflection fluorescence microscope (TIRFM). 80 bp Widom 601 DNA is tethered to the slide surface via biotin-streptavidin linkage in a microfluidic channel. After the tetrasome formation by CAF-1, QD-labeled FACTs (QD-FACTs) with H2A-H2B are imaged using TIRFM. (**B-D**). Quantification of single-molecule fluorescence imaging assays for FACT dissociation by (B) full-length Abo1, (C) Abo1ΔN, and (D) Abo1_W345A_. Each experimental schematic is presented at the right. The leftover number of QD-FACT/H2A-H2B on tetrasome is normalized by the leftover number in the absence of any Abo1 constructs. The experiment was repeated three times. For each quantification, 123, 163, and 127 molecules are analyzed for (B), (C), and (D) respectively, and error bars represent standard deviations in triplicate. (**E**) Quantification of single-molecule fluorescence imaging assay for FACT dissociation from tailless tetrasome by full-length Abo1. The leftover number of QD-FACT/H2A-H2B on tailless tetrasome is normalized by the leftover number in the absence of full-length Abo1. The experiment was repeated three times. For each quantification184 molecules are analyzed, and error bars represent the standard deviations in triplicate. Histone tail binding is required for FACT dissociation by Abo1.

We noticed that the QD signal decreased over time without Abo1 indicating that QD-FACT/H2A-H2B spontaneously dissociate from the tetrasome (Figure S5A), which hinders us from examining the FACT unloading activity by Abo1. We considered several other factors that could cause the reduction of QD signal without Abo1. The reduction of QD signals might result from the loss of surface-DNA binding, tetrasome disassembly, and/or the detachment of QD itself. We have found that tetrasome spontaneously dissembles from DNA (Figure S5B), suggesting that the reduction of the QD-FACT signal is due to the tetrasome instability over time. Therefore, in subsequent experiments, we normalized the amount of QD-FACT dissociation with that of spontaneous dissociation. With this setting, we examined whether Abo1 can affect the unloading of FACT from the tetrasome by monitoring the amount of QD signal. We have found that the number of QD-FACT was significantly reduced in the presence of Abo1 and ATP, indicating that Abo1 facilitates the dissociation of the FACT in an ATP-dependent manner (Figure 4B).

We have also investigated the possibility that Abo1 may influence the FACT loading onto tetrasome. To do this, we have pre-incubated QD-FACT/H2A-H2B with Abo1 before injecting the mix into a flowcell where tetrasome was immobilized. We have quantified the amount of QD to examine the enhancement of FACT loading by Abo1. Our experiments revealed that Abo1 does not increase FACT loading on the tetrasome (Figure S6), suggesting that Abo1 facilitates FACT unloading from chromatin rather than loading.

To further investigate whether the FACT dissociation depends on the physical binding of Abo1 and FACT, we measured FACT dissociation activity of Abo1ΔN, which has normal ATPase activity (Figure S7A) but lacks the FACT binding domain and is unable to form complexes with FACT (Figure 2B). Our results showed that the QD-FACT level did not change even when Abo1ΔN was added with ATP. These data indicate that FACT dissociation activity is mediated by the direct interaction between Abo1 and FACT (Figure 4C).

### Abo1 binding to histone tail via the central pore is required for FACT dissociation activity

Abo1 forms a hexamer with a central pore. We have previously shown that Abo1 interacts with histone tail by its central pore, and this interaction is essential for Abo1’s histone-loading activity (26). To investigate whether histone tail binding via the central pore is also necessary for Abo1-mediated FACT unloading, we generated Abo1 pore mutants (W345A or E385A), which retain normal ATPase activity and the ability to bind FACTΔC (Figure S7A and S7B). The single molecule analysis with Abo1 pore mutants showed that loss of QD-FACT from tetrasome was not accelerated, indicating that histone tail holding by the central pore is critical for FACT unloading activity of Abo1 (Figure 4D and S7C). We further examined the requirement of histone tail binding FACT dissociation activity using tailless histones. We generated the tailless tetrasome using tailless-H3/H4 with CAF-1 (Figure S7D and S7E) and measured the Abo1 FACT unloading activity. We found that Abo1 cannot dissociate FACT from tailless tetrasome, indicating that the histone H3-H4 tails are critical for Abo1 to dissociate FACT (Figure 4E). Collectively, the interaction between the central pore of Abo1 and histone tails is required for Abo1-mediated FACT dissociation.

### Abo1 dissociates only FACT from the FACT-histone complex on DNA

Our data shows that Abo1 facilitates the dissociation of FACT in a complex with H2A/H2B from H3/H4 tetrasome. We wondered what is the remaining component on DNA after FACT dissociation. We then aimed to examine the residual products following FACT dissociation. Abo1 interacts with histone tails upon FACT dissociation, potentially causing destabilization of the tetrasome structure. We investigated whether FACT dissociation coincides with tetrasome disassembly using electrophoretic mobility shift assay (EMSA). We pre-incubated FACT with H2A-H2B and loaded pre-formed FACT/H2A/H2B on tetrasome consisting of SYBR-Gold labeled 92 bp DNA and H3/_Cy5_H4 to load FACT on the tetrasome (Figure 5A, lane 3) (5,11). When Abo1 was added into the FACT-loaded tetrasome, the migration of FACT/H2A/H2B/tetrasome complex was retarded, indicating Abo1 binding to the FACT-loaded tetrasome (Figure 5A, lanes 4 and 7). In the presence of ATP, a new band between 300 to 400 bp emerged and became increasingly intense over time (30, 60, and 90 min) (Figure 5A, lanes 4-6), whereas it did not appear in the absence of ATP. The new band was observed in both signal channels for SYBR gold-labeled DNA and for Cy5-labeled histone H4, indicating that histones remained on DNA while Abo1 detached FACT from the tetrasome. We noticed that the gel mobility of the remaining product after FACT dissociation by Abo1 has substantially different to that of tetrasome alone, suggesting a possibility that the remaining product may contain H2A/H2B. To examine the existence of H2A/H2B in the remaining product, we conducted the EMSA using _Cy5_H2A/H2B. Our EMSA data show that H2A/H2B co-migrated with the produced tetrasome, indicating that Abo1 detached only FACT leaving H3-H4 and H2A-H2B on DNA (Figure 5B, lanes 4-6). To further determine the fate of histones after FACT dissociation, we performed a single molecule dissociation assay using H3/ _Cy5_H4 or _Cy5_H2A/H2B. Consistent with our EMSA analysis, the addition of FACT did not reduce the amount of H3/H4 from the DNA (Figure 5C). Regarding H2A-H2B, although there was a slight decrease in the level of _Cy5_H2A/H2B in the presence of ATP, the majority of H2A-H2B remained associated with the tetrasome even after FACT dissociation (Figure 5D). We speculated that the partial dissociation of H2A/H2B may result from the instability of H2A/H2B on the short 92 bp DNA.

**Figure 5.**
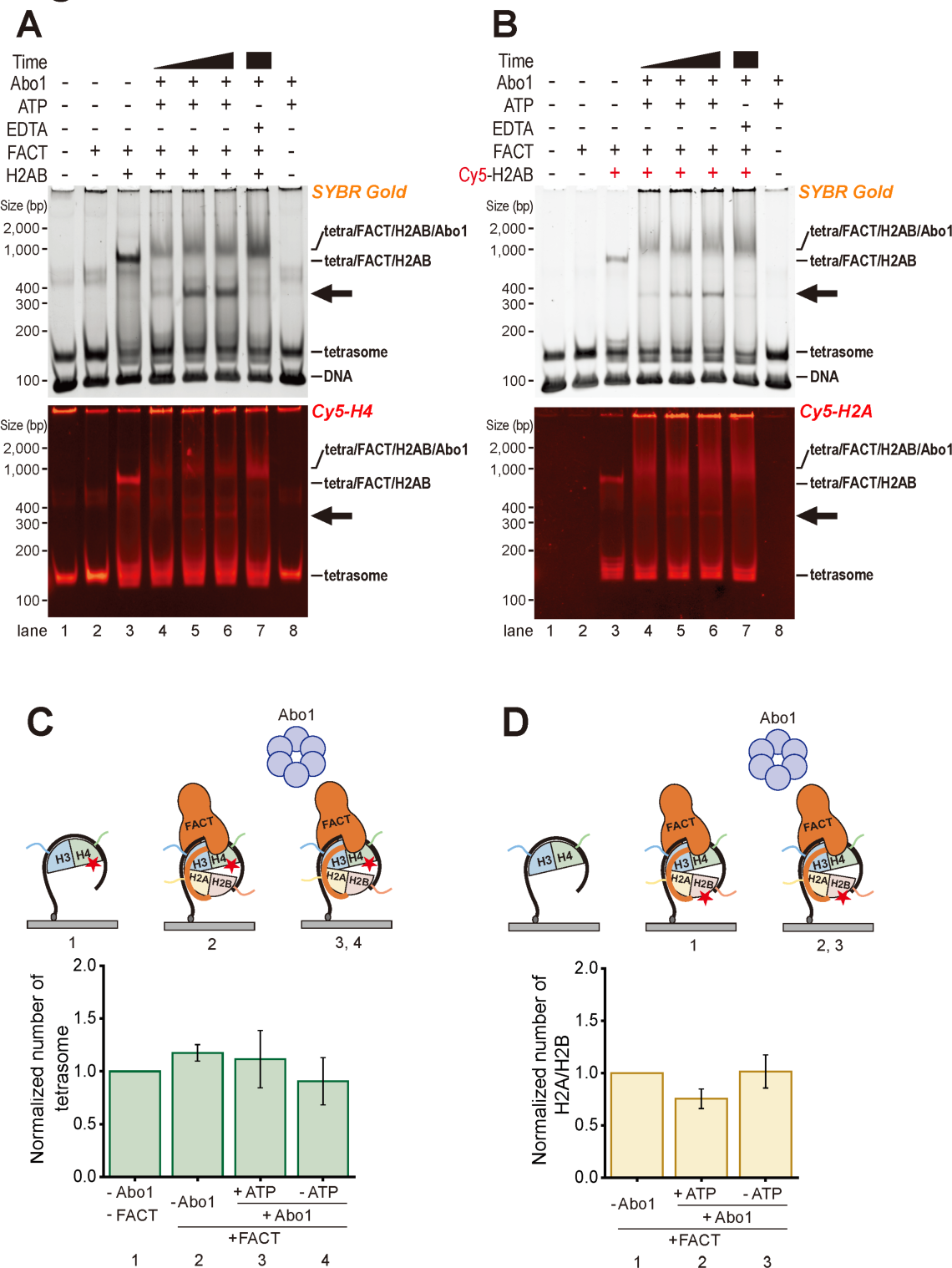
Abo1 does not disrupt tetrasome/H2A-H2B while dissociating FACT. (A) Analyzing the status of tetrasome after Abo1 dissociates FACT by Electrophoresis Mobility Shift Assay (EMSA). Tetrasome was assembled with 92 bp DNA and fluorescence-labeled H3-_Cy5_H4, where Cy5-maleimide was labeled on H4_T71C_. 100 nM tetrasome was incubated with 100 nM FACT/H2A-H2B for 30 min at RT (lane 3). 100 nM Abo1 and 1 mM of ATP were supplemented to tetrasome/FACT/H2A-H2B mixture for 30, 60, and 90 min (lane 4, 5, 6). 4 mM EDTA was added to the tetrasome/FACT/H2A-H2B to deplete residual ATP (lane 7). The samples were analyzed by 5% TBE native PAGE. Cy5-H4 was detected at 695nm, and DNA was stained with SYBR Gold. (B) H2A/H2B remains as tetrasome/H2A-H2B complex after Abo1 dissociates FACT. EMSA analysis with fluorescence-labeled _Cy5_H2A-H2B, where Cy5-maleimide was labeled on H2A_T120C_. The same condition was used in Figure 5A. (**C**) The fate of H3-H4 was examined by single-molecule analysis. The number of H3-_Cy5_H4, after the FACT dissociation in the presence of H2A-H2B. The experiment was repeated three times. 317 molecules were analyzed for each case and error bars represent the standard deviations in triplicate. Each number is normalized by the number in the absence of FACT and Abo1. (**D**) The fate of H2A-H2B was examined by single-molecule analysis. The number of _Cy5_H2A-H2B after the FACT dissociation. The experiment was repeated three times. 1055 molecules were analyzed for each case and error bars represent the standard deviations in triplicate. Each number is normalized by the number in the presence of FACT alone.

Lastly, we sought to determine whether Abo1 remains bound to tetrasome after the FACT dissociates by monitoring the level of Abo1. To visualize Abo1, we generated a 3×FLAG-Abo1 construct and fluorescently labeled it using anti-flag QD. We then compared the amounts of Abo1 under different flowcell conditions; with FACT/hexasome, tetrasome, or empty control (nonspecific binding to the surface). Our results showed that the level of Abo1 after the FACT dissociation reaction is similar to that of the empty control, although it appeared to be slightly reduced rather than exhibiting nonspecific surface binding. This suggests that Abo1 did not remain on the tetrasome after FACT removal (Figure S8). These data imply that Abo1 is also dissociated from the DNA-histone complex during FACT dissociation.

### The Abo1 N-terminal domain plays a crucial role in heterochromatin propagation

FACT plays critical role in heterochromatin silencing. Based on our finding that Abo1functions with FACT, we wondered how the Abo1-mediated FACT mobilization affects chromatin dynamics *in vivo* and decided to explore the effect of compromised FACT mobilization on heterochromatin stability. The fission yeast genome contains large blocks of heterochromatin defined by H3 lysine-9 methylation (H3K9me) at centromeres, telomeres, and silent mating type (*mat*) region (45). Heterochromatin assembly at the silent *mat* interval is initiated at the centromere homologous repeat element *cenH* (Figure 6A, cartoon), where RNAi machinery recruits H3 lysine-9 methyltransferase Clr4, the Suv39h homologue (46). The spreading of heterochromatin beyond the initiating *cenH* region and heritable propagation of repressive heterochromatic state requires critical density of trimethylated H3K9 (H3K9me3) (47,48), which serves to recruit Clr4 via its chromodomain to promote further deposition of H3K9me in surrounding regions (49). This read-write activity of Clr4 requires stabilization of nucleosome by chromatin modifiers that suppress turnover of histones to preserve high H3K9me3 density (10,50). The FACT complex associates with the central heterochromatin component, the H3K9me binding protein HP1, and promotes heterochromatin propagation by preventing loss of methylated nucleosome through histone turnover (8).

**Figure 6.**
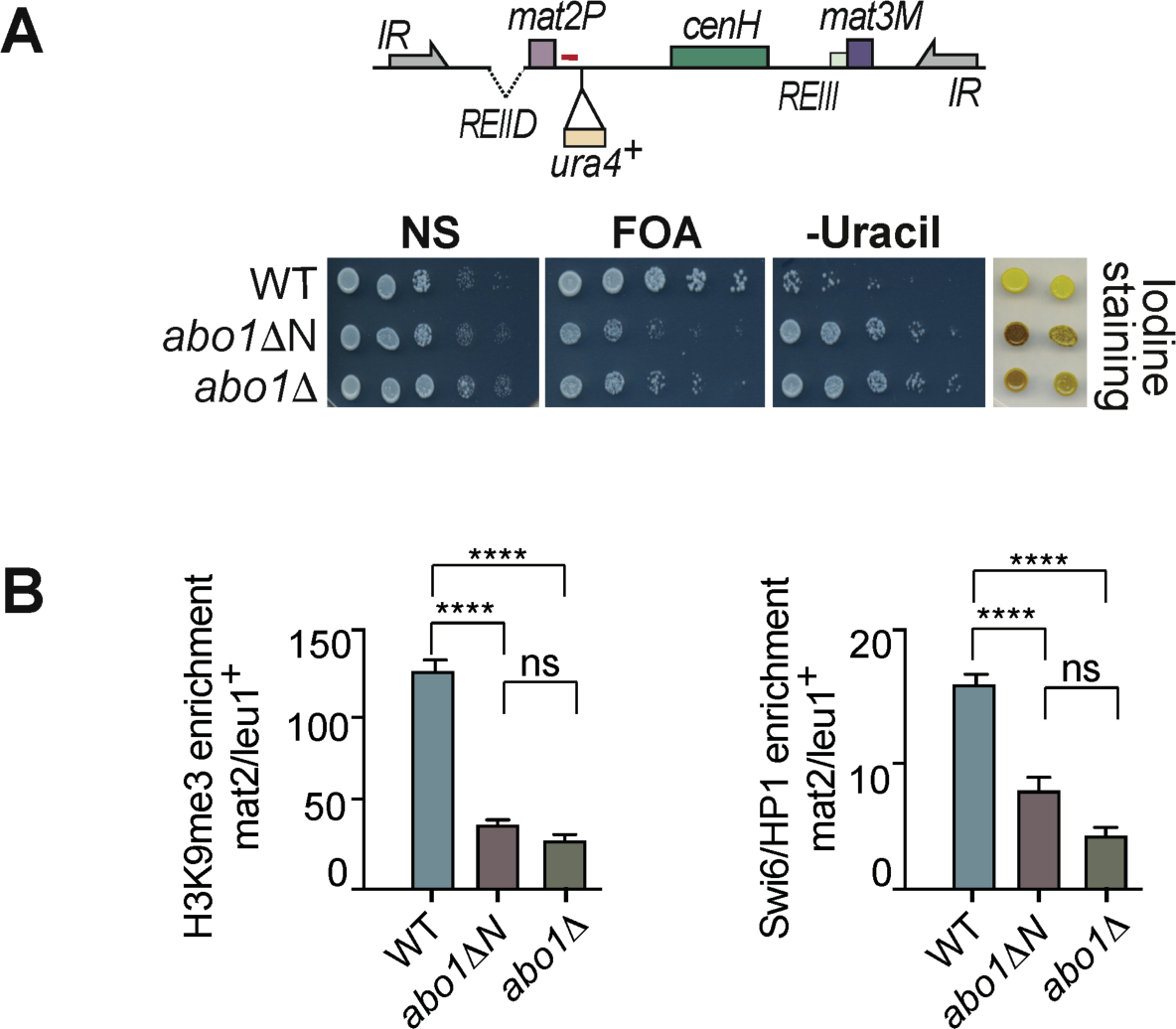
Abo1 N-terminal domains are required for heterochromatin silencing. (**A**) Abo1 N-terminal truncation impairs heterochromatin silencing. Serial dilution analysis of *mat2P::ura4^+^* heterochromatic repression. Serial dilutions of *WT* and *abo1* strains were spotted on non-selective (NS), uracil depleted (-URA) and *ura4^+^* counter selective FOA containing medium (+FOA). Cells grown on pombe minimal medium (PMG) were stain with iodine vapor to detect aberrant meiosis. (**B**) ChIP qPCR analysis of H3K9me3 and Swi6/HP1-immunoprecipitated DNA at *mat* region. Mean fold enrichments normalized to *leu1^+^* locus in indicated strains are plotted with bars indicating standard error of the mean of two experiments. The red bar in cartoon of panel A marks region analyzed. pValues of pair-wise comparison obtained by variance analysis are indicated above the columns; P>0.05: ns, P<0.0001: ****.

To investigate the role of FACT mobilization by Abo1 in heterochromatin propagation, we examined heterochromatin at the silent *mat* region in cells expressing Abo1 that is missing the N-terminal region required for FACT dissociation. We employed a highly sensitive reporter that monitored heterochromatin propagation maintained by pre-existing methylation. Deletion of local *REII* silencer at *mat2* cassette (*REIIΔ mat2::ura4^+^;* Figure 6A, cartoon) renders *mat2* repression dependent solely on heterochromatin. In non-switching *mat1M* haploid cells compromised maintenance of H3K9me3 density derepresses *mat2P* cassette and *mat2-*proximal *ura4^+^* reporter and initiates aberrant meiosis process that results in spore formation detected by brown staining with iodine vapor. Cells lacking Abo1 (*abo1Δ)* or expressing the N-terminally truncated Abo1 *(abo1ΔN*) derepressed *ura4^+^* reporter (Figure 6A), as indicated by enhanced growth on medium lacking uracil (-URA) and suppressed survival on *ura4^+^* counter-selective medium containing 5-fluoroorotic acid (+FOA). Moreover, both *abo1Δ* and *abo1ΔN* cells stained brown when exposed to iodine vapors, demonstrating initiation of aberrant meiosis caused by *mat2P* derepression (Figure 6A). Derepression was linked to defective heterochromatin assembly, as indicated by a considerable reduction in the levels of both H3K9me3 and Swi6/HP1 localization at *mat2* in *abo1* mutant cells when compared to WT cells (Figure 6B).

Our analysis indicates that the NTR of Abo1, required for FACT mobilization, is also crucial for the epigenetic propagation of heterochromatin through the Clr4 read-write mechanism. This demonstrates the importance of Abo1 in maintaining proper heterochromatin formation and gene silencing in fission yeast cells.

## Discussion

In this paper, we dissected the direct interaction between Abo1 and FACT and investigated the role of Abo1 on the function of FACT, showing that Abo1 facilitates the dissociation of FACT from chromatin in an ATP-dependent manner (Figure 7). Abo1 maintains global nucleosome organization and is involved in heterochromatin silencing (25,28). In addition, Abo1 depletion affects the level of FACT component Spt16, and controls the turn-over of HIRA histone chaperone (24,28). These previous studies suggest diverse functions of Abo1 involving interaction with other proteins. We showed that Abo1 directly interacts with the N-terminal domain (NTD) of Spt16 using the FIN helix located at the N-terminal region (NTR). The NTR of Abo1 exhibits low sequence conservation among its homologues, suggesting that functional differences between Abo1 homologues may be linked to this N-terminal variability (27,29,30). The precise role of the NTR in Abo1’s function remains unclear, but it may serve as a domain for protein-protein interaction domain. Supporting this idea, human Abo1, ATAD2 is shown to interact with the Androgen receptor or E2F through its N-terminal region (23,51). Therefore, it is plausible that the NTR of Abo1 homologues plays a crucial role in protein-protein interactions. This includes interactions with histones and other binding partners, utilizing the charged region of the N-terminal domain. Based on our previous cryo-EM structure, the NTR of Abo1 is topologically located near to the bromodomains. The bromodomain together with the ATPase domain undergoes large conformational change upon ATP hydrolysis (26). However, Abo1 does not seem to interact with FACT in an ATP-dependent manner, indicating that the N-terminal binding site is exposed regardless of the structural change of Abo1. The Spt16-NTD contains an aminopeptidase fold with no catalytic activity. Instead of Spt16-NTD having an enzymatic activity, it was shown to serve as an interaction site for various proteins, including histone H3-H4, CHD1, Tof1, and Swi6 (13,15,18,52–54). Our data also demonstrates that Spt16-NTD is used for interacting with Abo1 FIN helix. The interaction with Abo1 via Spt16-NTD seems to depend on the FACT conformation. A previous work showed that the C-terminal regions of FACT control FACT conformation (36). We have observed that Abo1 interacts with FACT lacking C-terminal regions, but not with full-length FACT. Additionally, the full-length FACT becomes competent to bind to Abo1 when its C-terminus is engaged by H2A/H2B. These data indicate that FACT C-terminal region masks the Abo1 binding interface of Spt16-NTD and that H2A/H2B binding to FACT releases Spt16-NTD for Abo1 interaction. It can be speculated that the C-terminal region of FACT prevents FACT from nonspecific interaction with DNA or other interaction partners in absence of H2A/H2B dimer. This regulation could facilitate precise control of FACT’s involvement in cellular processes. Previously we have shown that Abo1 loaded histones on DNA in an ATP-dependent manner. Interestingly, the binding of histone tails by Abo1 pore residues is necessary for both histone loading, and for Abo1-mediated FACT dissociation. Many AAA+ proteins bind the substrate using a central channel, pulling the substrate to induce unfolding activity (55). In this manner, Yta7 is known to extract H3 from chromatin. However, Abo1, by anchoring the histone tail of the extended tetrasome, does not disrupt the tetrasome structure. Instead, it appears to dissociate FACT while leaving the chromatin intact selectively. At this moment, it is uncertain whether the two functions of Abo1, histone loading activity and FACT dissociation activity, are separate or interconnected.

**Figure 7.**
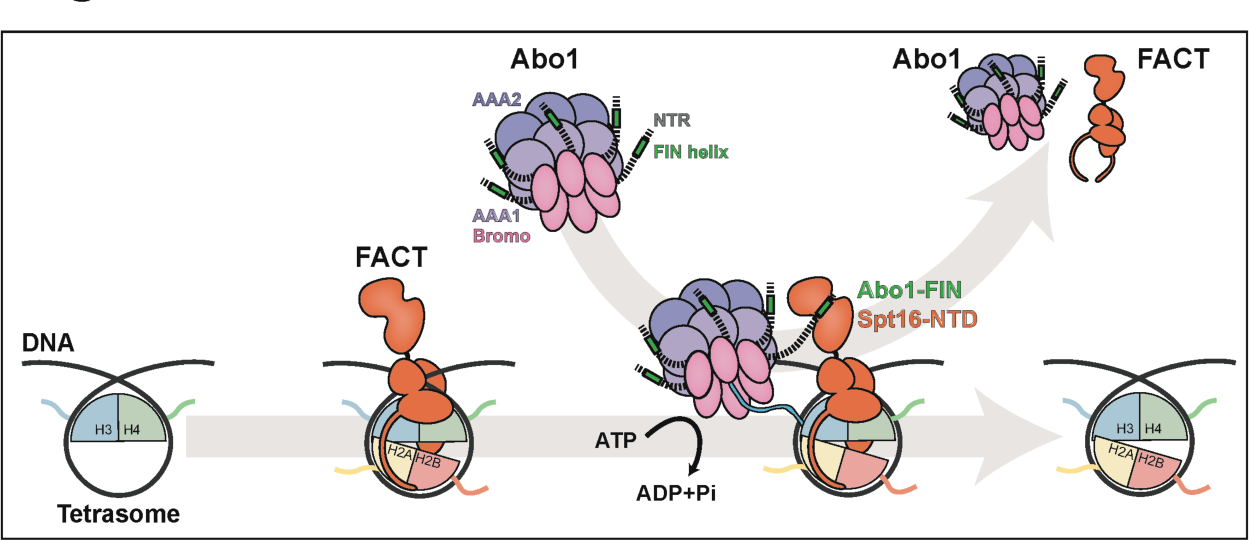
A schematic model of Abo1-mediated FACT dissociation. Abo1 has an N-terminal region (NTR, dashed line) that includes a FACT interacting helix (FIN, green), two ATPase domains (AAA1, 2, purple), and a Bromodomain (Bromo, pink). FACT binds to an opened chromatin-like tetrasome, with Spt16-NTD protruding from chromatin. Abo1 interacts with Spt16-NTD through its FIN helix while simultaneously engaging the histone H3 tail (light blue) with its pore residues. Utilizing ATP energy, Abo1 dissociates FACT and leaves chromatin with DNA, H2A-H2B, and H3-H4.

In this study, we discovered another role of Abo1 ATPase activity in disengaging FACT from chromatin in addition to histone loading activity. Considering the high abundance of FACT in cells, the primary function of Abo1-mediated FACT release may be rather coordination of chromatin remodeling in processes such as transcription or heterochromatin silencing than FACT recycling. Although the recycling mechanism of the FACT histone chaperone complex remains poorly understood, our finding offers initial insights into the ATP-dependent FACT recycling facilitated by Abo1. Our work presents an example of histone chaperone effecting chromatin conformation by controlling another chaperone protein rather than histone.

## Data Availability

The data underlying this article are available in the article and in its online supplementary material.

## Supplementary Data

Supplementary Data are available at NAR online.

## Author contributions

J.J., Y.K., J.L. and J.S. conceived the study, designed and performed biochemical and single-molecule experiments. M.Z. designed and performed all in vivo experiments. S.G. designed and supervised all in vivo experiments. C.C. conceptualized the study and edited the manuscript. J.Y.L. and J.S. supervised experiments. All reviewed the data, wrote and revised the manuscript.

## Acknowledgments

We thank Song lab members for helpful discussion, and Yumi Shin and Eunhee Seong for their technical assistance. We thank Seong-Woo Kim of KAIST for the helpful discussion about AlphaFold prediction.

## Funding

National Research Foundation of Korea (RS-2024-00333346, RS-2023-00266300 to J.S.), The Samsung Science and Technology Foundation (SSTF-BA1901-13 to J.Y.L.)

## Conflict of Interest

J.S. is a CTO of Epinogen.

---

**Figure S1.**
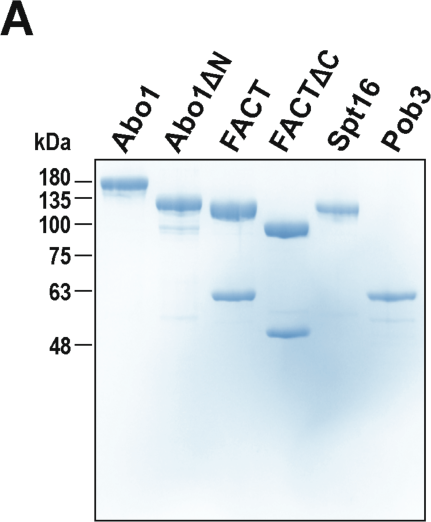
Purified Abo1 and FACT proteins. Purified Abo1, Abo1ΔN, FACT, FACTΔC, Spt16, and Pob3 were subjected to analysis using 10% SDS-PAGE.

**Figure S2.**
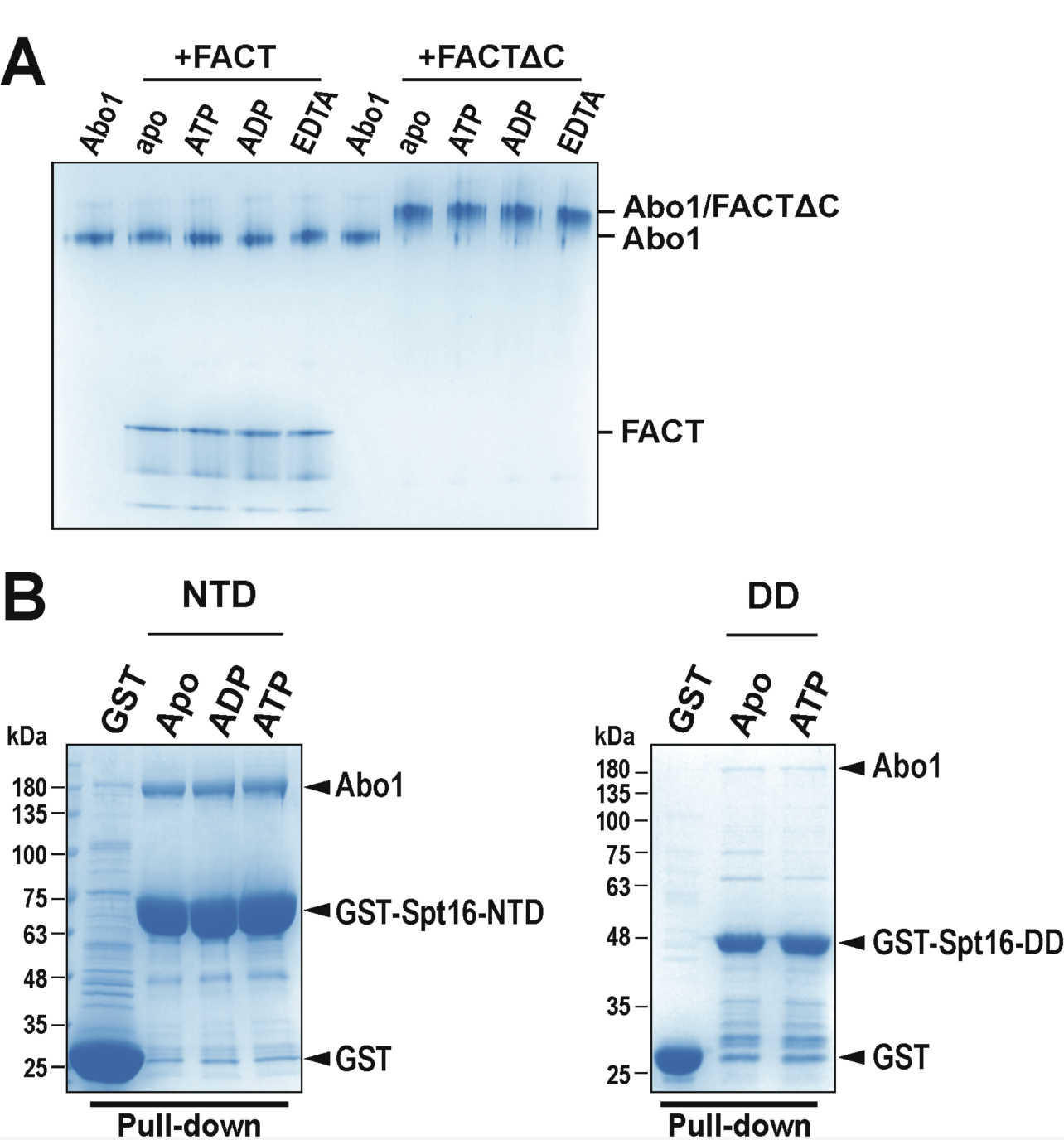
Abo1 does not require ATP for FACT binding. (**A**) 100 nM Abo1 was incubated with 400 nM FACT or FACTΔC in the presence of 1 mM ATP, 1 mM ADP, or 4 mM EDTA for 30 min at RT. The complex formation was analyzed by 3-12% Bis-Tris native PAGE. (**B**) Abo1 binds to Spt16-NTD (left panel) or Spt16-DD (right panel) in an ATP-independent manner. The GST pull-down assays were conducted as described in Figure 2C in the presence of 1 mM of ATP or ADP. The bindings were analyzed by 10% SDS PAGE.

**Figure S3.**
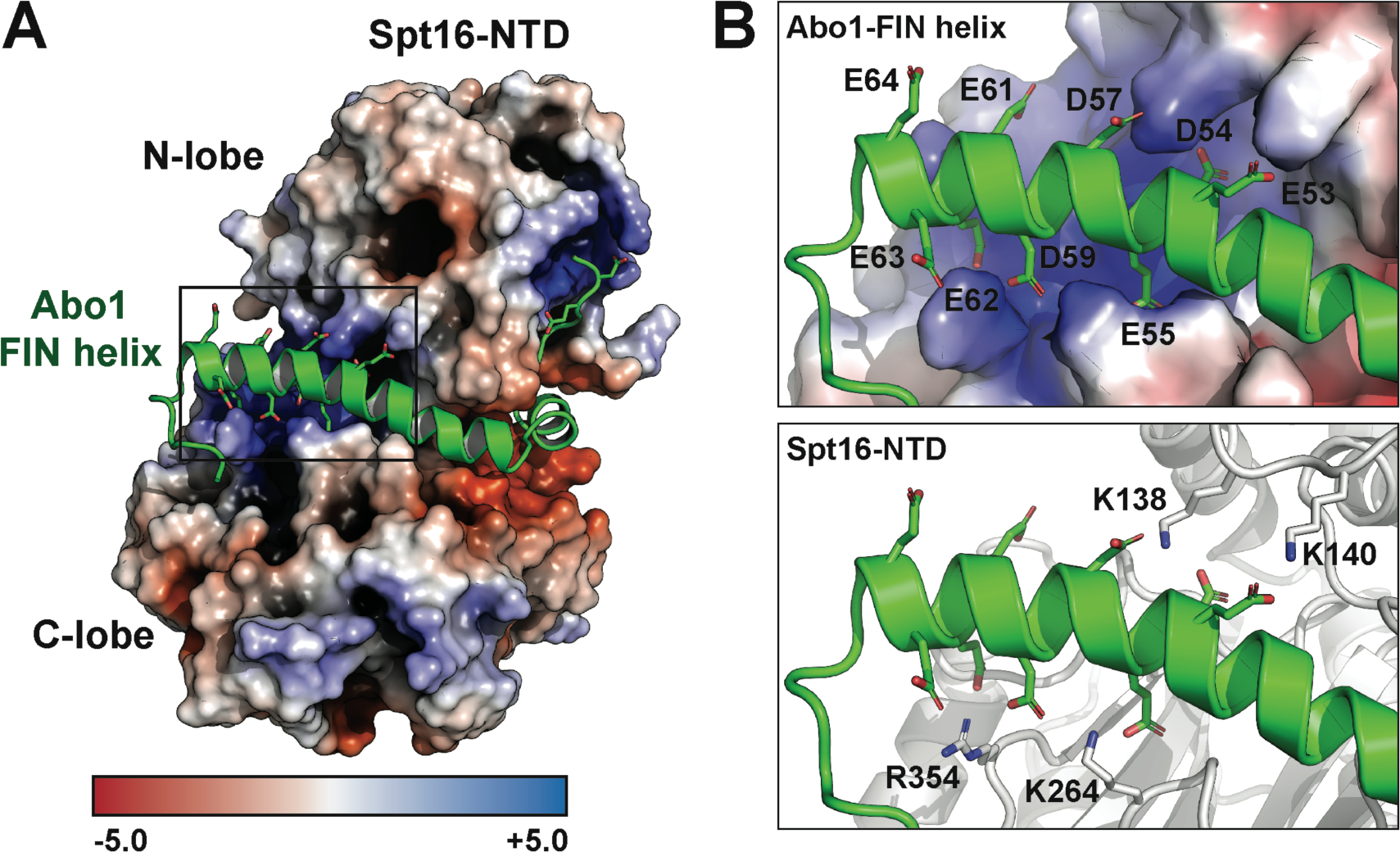
FIN helix binds to the basic patch of Spt16-NTD. (A) The interaction between the acidic residues of the FIN helix and a basic patch of Spt16-NTD. The Spt16-NTD is shown as the electrostatic potential surface map. (**B**) Detail interactions between FIN helix and Spt16-NTD. (Top) Acidic residues of the FIN helix are bound to the basic patch of Spt16-NTD. E53, D54, E55, D57, D59, E61, E62, E63, and E64 of FIN helix were visualized. The blue region represented the basic patch of Spt16-NTD. (Bottom) Basic residues of Spt16-NTD interact with the FIN helix. K138, K140, K264, and R354 were visualized.

**Figure S4.**
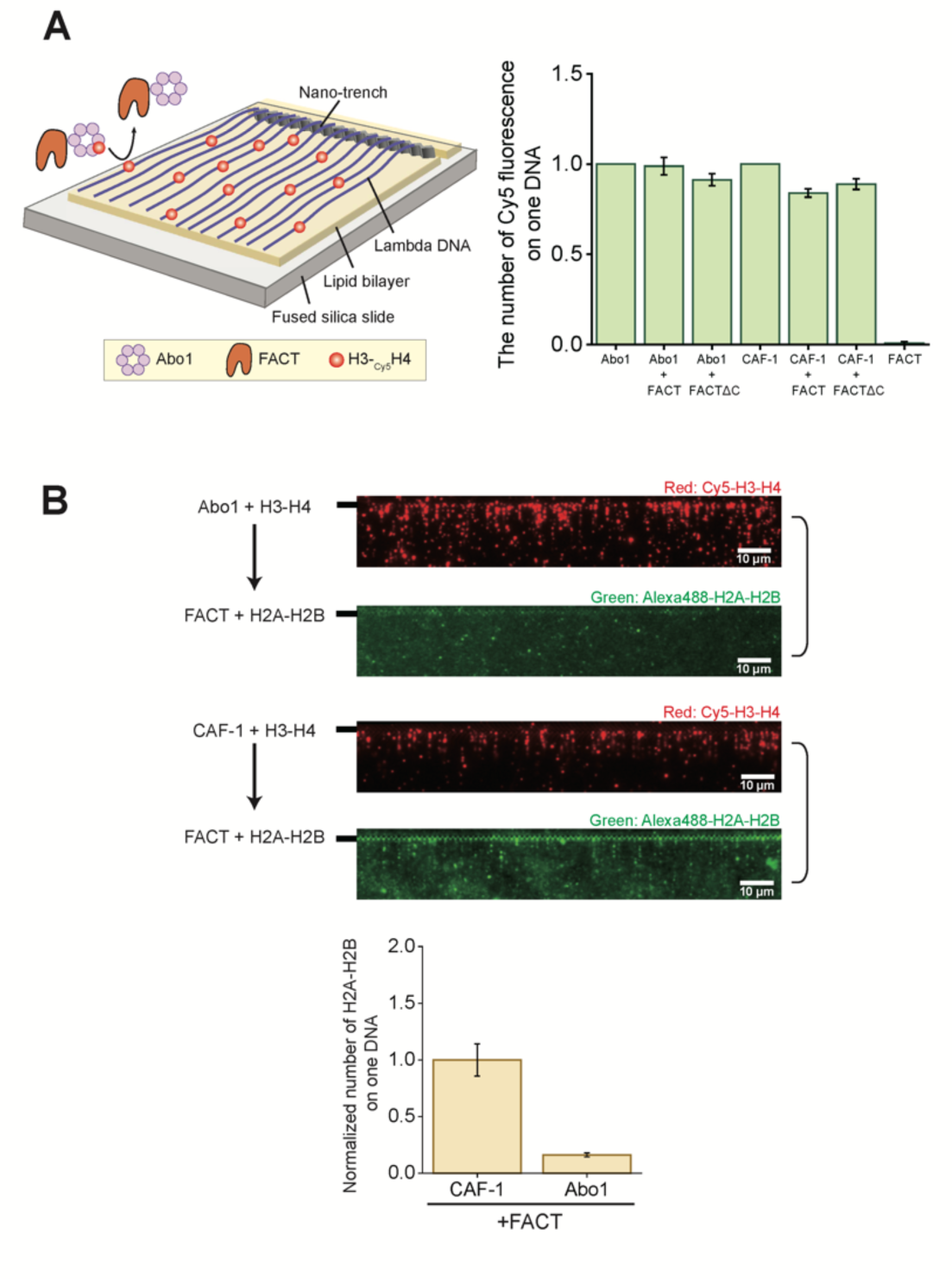
Abo1 and FACT do not cooperate in nucleosome assembly. (**A**) Schematic of single-tethered DNA curtain assay (left panel). The effect of FACT on H3-H4 loading activity (right panel). H3-_Cy5_H4 dimers are deposited onto DNA by Abo1 or CAF-1 with or without FACT (or FACTΔC). H3-H4 deposition onto DNA by Abo1 or CAF-1 with or without FACT (or FACTΔC) is quantified. The data in the conditions in the presence of FACT are normalized with the value of Abo1 or CAF-1 alone. The experiment was repeated three times. For each quantification, average 97.7 DNA molecules are analyzed. Error bars represent standard deviation in triplicate. (**B**) DNA curtain images for the formation of nucleosome. _Alexa488_H2A-H2B dimers (green) and FACT are added to DNA molecules, on which H3-_Cy5_H4 dimers (red) are pre-loaded by Abo1ΔN (top panel). _Alexa488_H2A-H2B dimers (green) and FACT are added to DNA molecules, on which H3-_Cy5_H4 dimers (red) are pre-loaded by CAF-1 (middle panel). FACT can assemble H2A-H2B into the tetrasome formed by CAF-1. The black bars at the left indicate the barriers. Quantification data for the formation of nucleosomes (bottom panel). The experiment was repeated three times. For each quantification, 95 DNA molecules are analyzed. Error bars represent standard deviation in triplicate.

**Figure S5.**
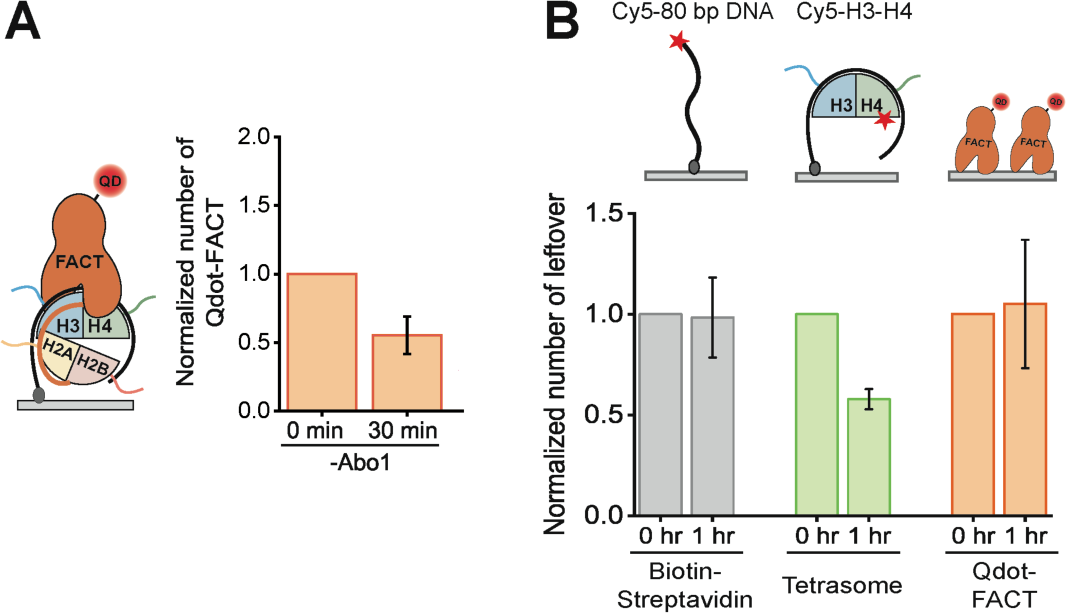
Spontaneous FACT dissociation from tetrasome instability. (**A**) Schematic of QD-FACT/H2A-H2B binding to a tetrasome on the surface (left panel). Quantification of the spontaneous decrease of QD-FACT (right panel). The leftover number of QD-FACT at a tetrasome in the presence of H2A-H2B. The experiment was repeated three times. For each quantification, 231 molecules are analyzed. Error bars represent standard deviation in triplicate. (**B**) Control experiments for the single-molecule fluorescence imaging assay for FACT dissociation. The normalized leftover number of DNA from the surface (grey), tetrasome (green), and QDs conjugated with FACT (orange) during the reaction (1 hour). In 1 hour, the leftover fraction of tetrasome was approximately 58%, indicating that tetrasomes are spontaneously disassembled. The schematic of each control experiment is shown above. The experiment was repeated three times. For each quantification, 83, 64, and 222 molecules are analyzed, tetrasomes, and QD-FACT respectively, and error bars represent standard deviations in triplicate.

**Figure S6.**
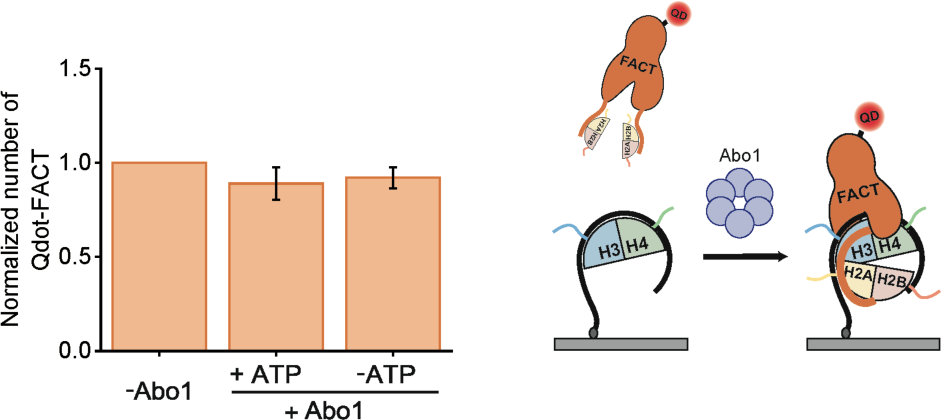
Single-molecule assay for FACT loading by Abo1. The number of QD-FACT/H2A-H2B that are left on tetrasome in the presence of full-length Abo1. The leftover number is normalized by the number in the absence of full-length Abo1. For all quantified data, more than 100 molecules were analyzed and error bars represent the standard deviation in triplicate. The schematic for FACT loading by Abo1 is shown at the right of the quantified data. The experiment was repeated three times. For each quantification, 148 molecules are analyzed and error bars represent standard deviations in triplicate.

**Figure S7.**
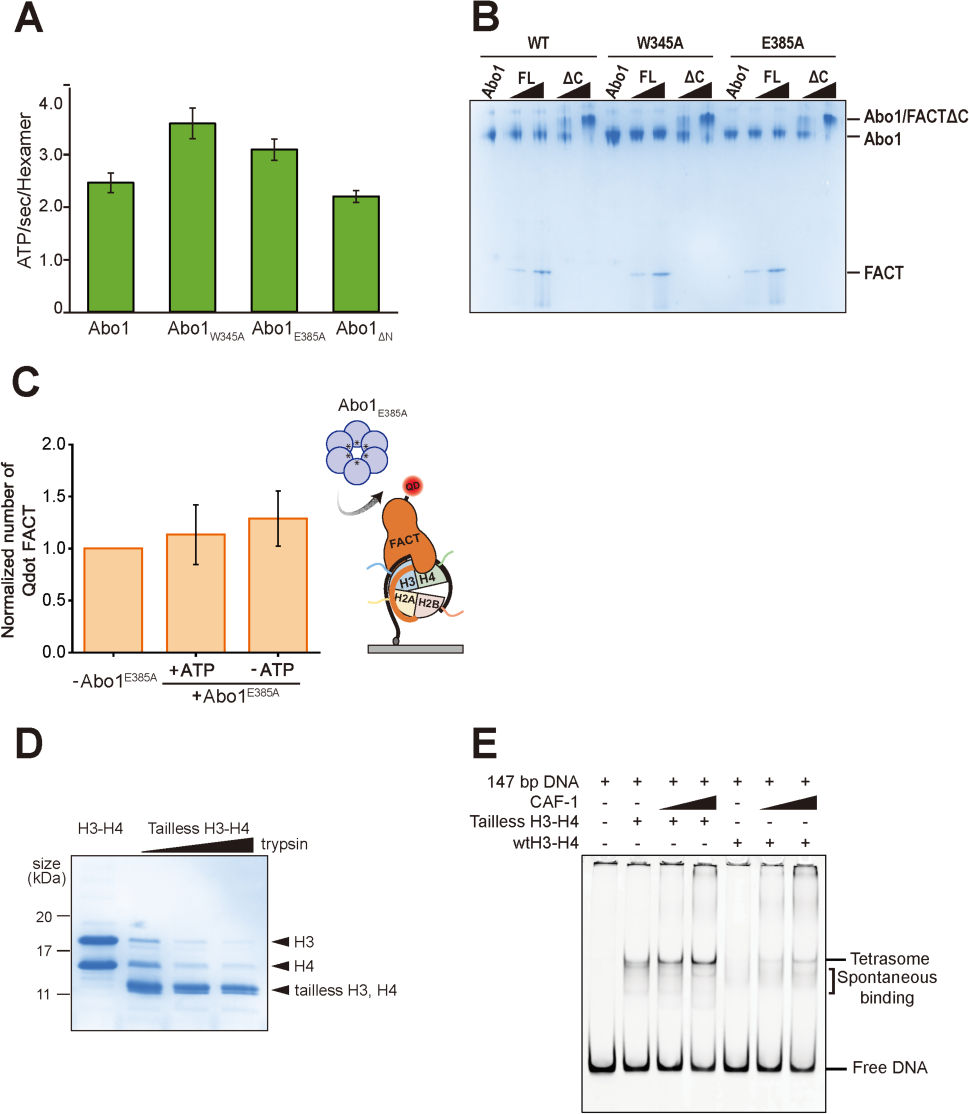
Histone tail binding by Abo1 pore residues is required for FACT dissociation. (**A**) ATPase activity of Abo1 and Abo1 mutants. Abo1, W345A, E385A, and Abo1 ΔN were tested for ATPase activity. Error bars represent the standard deviation for three experiments. (**B**) FACT binding affinity of Abo1 and Abo1 mutants. In the same way as Figure 1A, 100 nM of Abo1 was incubated with 100 or 400 nM of FACT (or FACTΔC) for 30 min at RT. The resulting proteins were separated using 3-12% Bis-Tris native PAGE and stained with Coomassie. (**C**) The normalized leftover number of QD-FACT by Abo1_E385A_. FACT is barely dissociated from a tetrasome by FLAbo1_E385A_. The experiment was repeated three times. 93 molecules are analyzed for each case. Error bars represent the standard deviations in triplicate. (**D**) Preparation of tailless H3-H4 by treatment trypsin. H3-H4 was cleaved by trypsin, with the treatment time increased from 10 to 30 minutes, and the cleavage was confirmed by 15% SDS-PAGE. (**E**) Tetrasome formation assay for CAF-1. 50 nM of Widom 601 (147 bp) was incubated with CAF-1 (0, 100, and 200 nM) pre-incubated with 50 nM of WT H3-H4 or tailless H3-H4. Tetrasome band becomes dominant as CAF-1 concentrations increase both with WT H3-H4 and tailless H3-H4.

**Figure S8.**
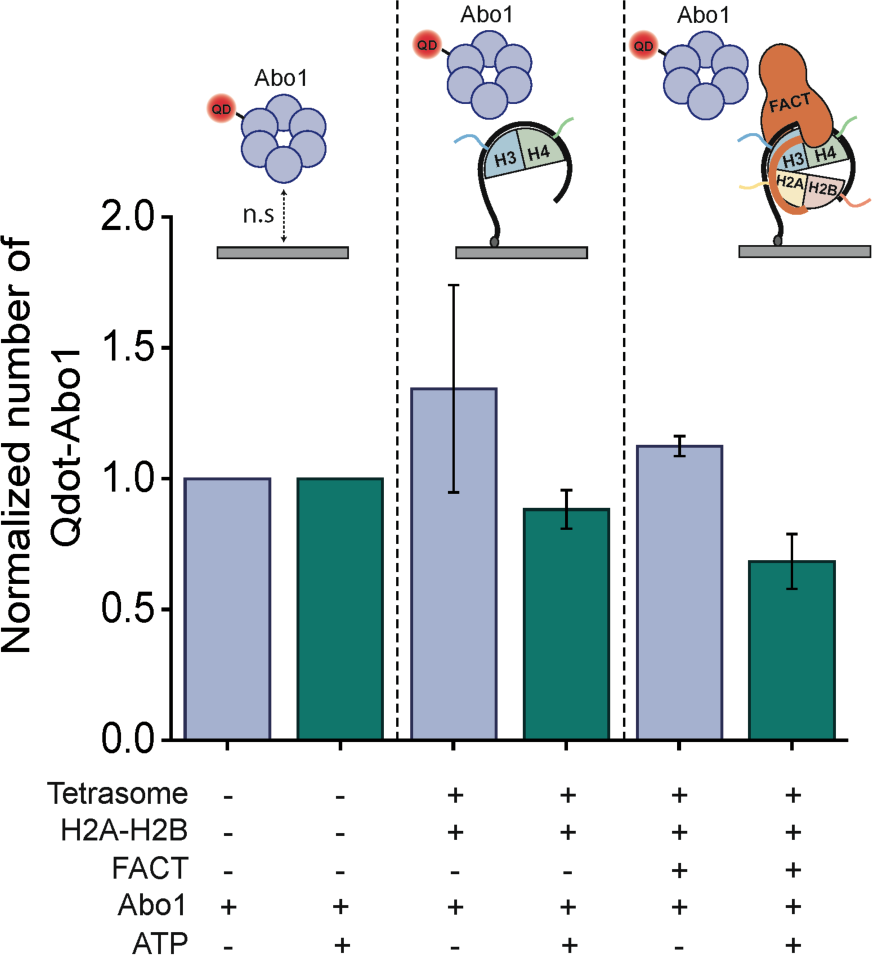
The fate of Abo1 after FACT dissociation. The normalized number of QD-FLAbo1s with the FACT dissociation in the absence or presence of ATP. The experiment was repeated three times. 32 molecules are analyzed for each case and error bars represent standard deviations in triplicate. The schematic for each case is shown at the top of the quantified data.

**Table S1:**
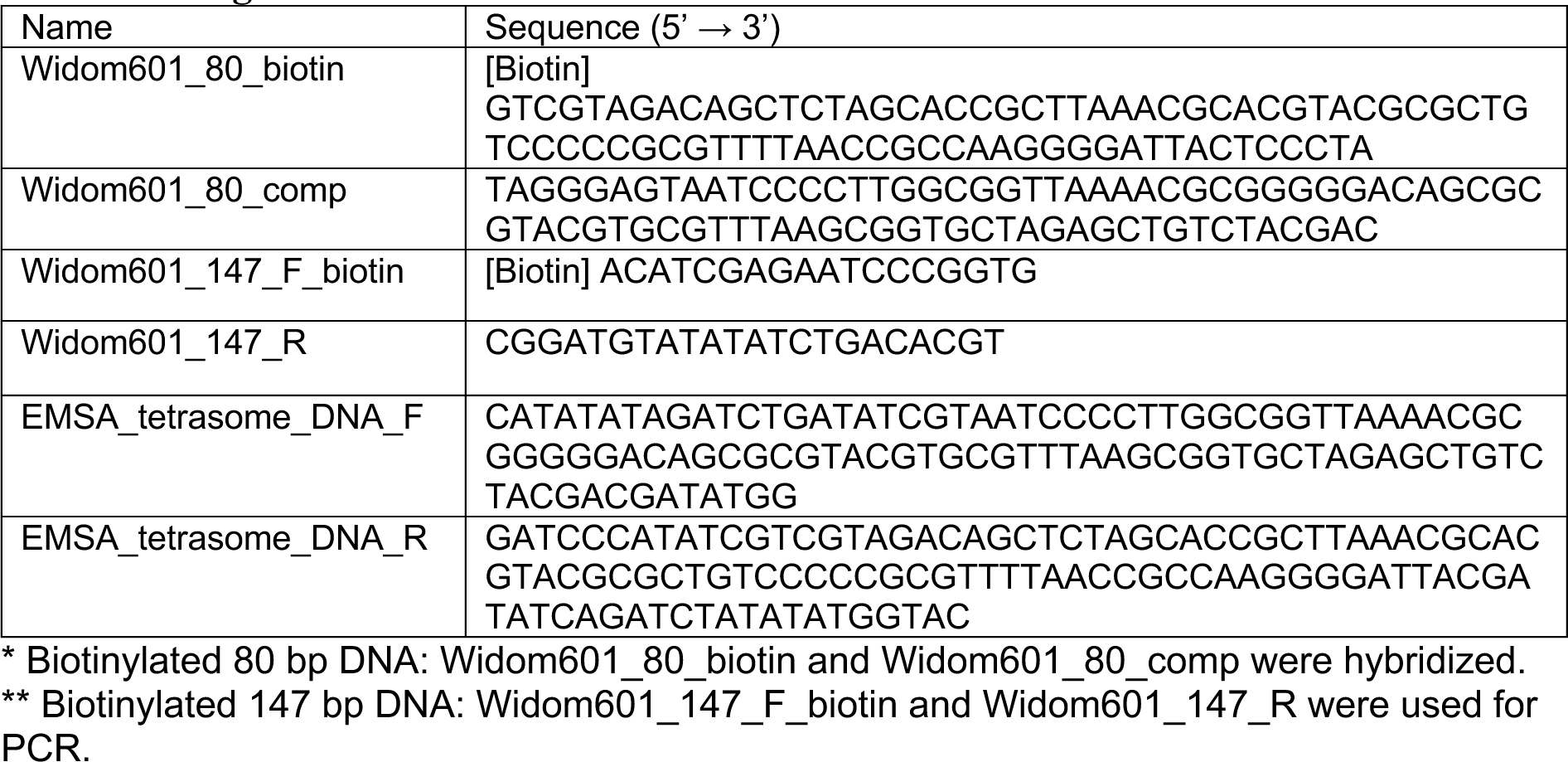
Oligomer List.

**Table S2:**
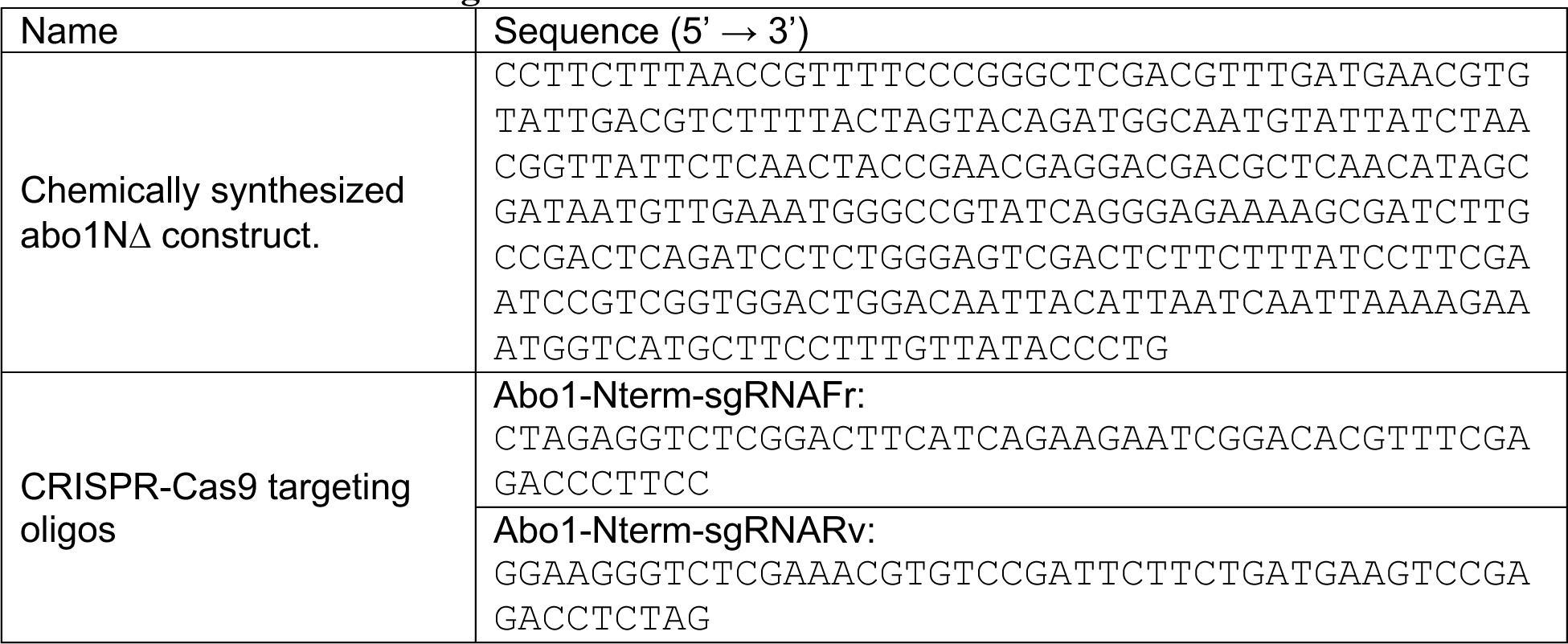
Construction Oligos.

**Table S3:**
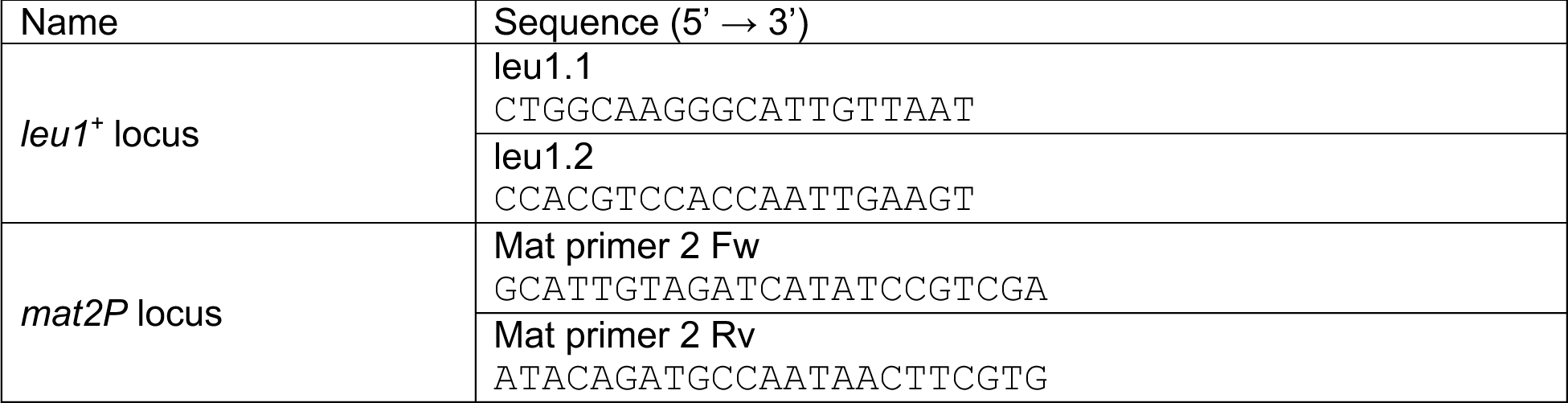
qPCR Oligos.

